# Virtual lineages: in silico lineage tracing from single-cell and spatial transcriptomes

**DOI:** 10.64898/2026.10.19.760045

**Authors:** Yongjie Lin, Yihan Lin

## Abstract

Experimental lineage tracing reveals histories and fates of individual cells but is costly and often impractical, particularly in primary human tissues. Gene expression differences that persist across cell divisions provide a basis for inferring ancestry from more accessible single cell transcriptomes, yet accurate lineage reconstruction remains challenging. Here, we introduce isCORAL, an in silico lineage-tracing framework that reconstructs clonal relationships and maps cell-fate dynamics from transcriptome snapshots alone. By identifying memory genes with heritable expression differences, isCORAL infers probabilistic clonal relationships and uses them to construct clone-oriented embeddings and dynamic cell-fate maps. Across diverse lineage-tracing datasets, isCORAL markedly outperforms tested alternatives in both memory-gene identification and lineage reconstruction. We use isCORAL to resolve early fate biases among transcriptionally similar hematopoietic stem cells and predict distinct developmental routes to shared outcomes. We extend isCORAL to spatial transcriptomics, validating inferred clonal relationships against an experimentally recorded lineage tree in mouse tumors and inferring early epiblast and amnion dynamics in a human embryo. Together, isCORAL enables lineage-resolved studies of development and disease from standard single cell and spatial transcriptomes, and provides an explicit representation of lineage memory for future virtual cell models.

## Introduction

Understanding how mammalian cells acquire their fates during development and disease progression is a central goal of cell biology^1^. Single cell omics has transformed our ability to characterize cell states and provides a foundation for predictive virtual cell models^2–4^, but most assays capture static snapshots^5,6^. Computational approaches, including pseudotime ordering^7,8^, RNA-velocity models^9–11^, dynamical systems models^12–14^ and optimal transport^15,16^, infer developmental trajectories and transition probabilities from these data. Yet the ancestry and future fates of individual cells remain difficult to infer reliably from transcriptomes alone^17,18^.

Experimental lineage tracing complements these inferences with an independent record of cellular ancestry, obtained by labeling individual cells and following their progeny^6,19,20^. Recent lineage-tracing technologies, from static lentiviral barcoding such as LARRY^17^ to highly multiplexed CRISPR-based genome editing such as DARLIN^21^ and DNA Typewriter^22^, have reshaped our understanding of cell dynamics across organisms, especially when combined with single-cell omics.

Despite their utility, lineage-tracing modalities remain far less accessible than large-scale scRNA-seq atlases. Prospective strategies generally require the experimental introduction of heritable barcodes, making them labor-intensive and largely inapplicable to intact human tissues in vivo. Retrospective strategies based on intrinsic mutations (e.g., somatic^23,24^ or mitochondrial mutations^25,26^) avoid introducing exogenous labels, but informative mutations are often sparse, stochastically acquired, and incompletely captured by standard single-cell assays^27^, limiting reconstruction accuracy and temporal resolution^27^, particularly over short developmental intervals.

These limitations raise the question of whether transcriptomic patterns contain enough information for short to medium timescale lineage reconstruction, as genomic^23^ and epigenomic modalities^28^ do over longer timescales. Addressing this question remains challenging because gene expression is noisy and changes with cell state^29^. Nevertheless, some gene-expression differences persist across cell divisions^30–33^. This persistence is termed transcriptional memory, and the genes displaying it are termed memory genes. Using experimental lineage labels, our CORAL study^32^ identified widespread transcriptional memory and showed that memory genes predict metastatic potential and lineage commitment. These inherited expression differences can help distinguish cells with different fate biases within the Waddington landscape^5,34^.

Thus, identifying transcriptional memory offers a route to lineage inference from standard single-cell data, motivating transcriptome-based *in silico* lineage-tracing methods. Existing approaches address different parts of this problem: lineage-informed velocity methods use measured genealogies to identify transcriptomic clocks^35^; GEMLI predicts clonally related cells from expression memory^33^; and Power-Seek estimates memory strength without explicitly reconstructing clone-level structure^36^. However, it remains highly challenging to reconstruct how clonally related cells are distributed across changing cell states with sufficient accuracy to support quantitative fate mapping.

Here we introduce isCORAL (in silico clone-oriented reconstruction of a landscape), an interpretable statistical framework for lineage reconstruction and quantitative cell-fate analysis from transcriptomes. Its clonal-inference module is unsupervised and requires no experimental lineage barcodes. First, isCORAL identifies memory genes and estimates, for each cell, a probabilistic distribution of clonally related cells, termed a “pseudoclone”. The downstream pCORAL module, adapted from our experimental lineage-analysis framework CORAL^32^, organizes cells according to the distributions of their inferred clonal relatives. This representation supports analysis of fate bias and developmental routes within standard single cell workflows. isCORAL outperformed tested alternatives in consistently recapitulating gene heritability and clonal structures across diverse cancer and developmental systems. Applied to hematopoietic lineage tracing^37^, mouse tumor spatial transcriptomics^38^ and human embryo spatial transcriptomics^39^, isCORAL successfully resolved early fate decisions and developmental trajectories that were obscured in conventional analyses. Together, these results show that the lineage information latent in ordinary transcriptomes can be recovered computationally, extending lineage-resolved analysis to the many settings where experimental tracing is impossible.

### Heritable memory genes define a feature space for in silico lineage tracing

The central challenge of in silico lineage tracing is that the transcriptome changes continuously with cell state: transcriptomes of different clones overlap substantially in high-dimensional expression space, making clones difficult to distinguish^29^ (**Figure 1A**). However, recent studies across diverse biological systems found that certain genes can maintain their expression levels for several generations^30–32^ (spanning days to weeks), leading to high variance between clones relative to within them (**Figure 1B**). These observations suggest that such highly heritable memory genes encode lineage information more effectively than the global transcriptome. We therefore hypothesized that in silico lineage tracing could be realized if probabilistic clone inference were performed within a feature space defined by carefully selected memory genes.

**Figure 1.**
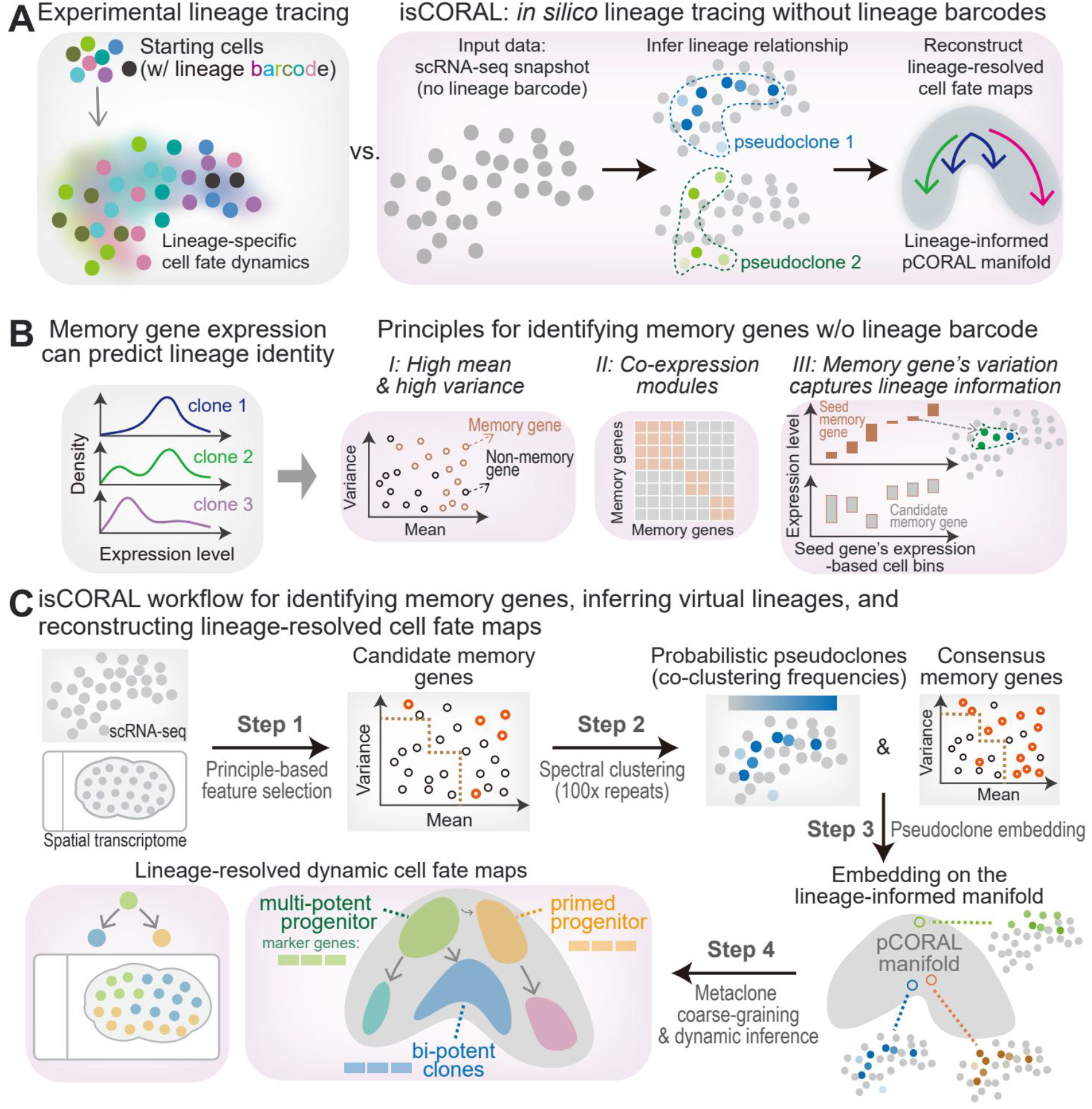
Heritable memory genes define a feature space for in silico lineage tracing. (A) Motivation and overview of isCORAL, an *in silico* lineage-tracing workflow. Experimental lineage barcodes connect starting cells to their descendants (left). isCORAL instead infers probabilistic pseudoclones from unbarcoded single-cell transcriptomic snapshots and uses them to construct a lineage-informed pCORAL manifold and reconstruct lineage-resolved cell-fate maps (right). (B) Principles for identifying memory genes without experimental lineage barcodes. Clonally persistent expression allows memory genes to distinguish lineage identities, as illustrated by the expression distributions at left. Three principles guide feature selection: (I) memory genes tend to have high expression and/or high variance; (II) co-expressed memory genes can be evaluated as modules; (III) expression bins defined by a seed memory gene can enrich cells with shared clonal ancestry, allowing other memory genes to be identified through variation across these bins, even when their relationship with the seed gene is not linear. (C) The isCORAL workflow in four steps. (1) Principle-guided feature selection identifies candidate genes with a piecewise mean–variance criterion. Non-negative matrix factorization (NMF) detects candidate modules; temporary spectral-clustering bins are used to evaluate the modules and retain key memory features. (2) Joint pseudoclone and memory-gene inference: spectral clustering on the selected key features yields a temporary pseudoclone partition and gene-wise omega-squared scores. Repeating feature selection and clustering 100 times yields cell-pair co-clustering frequencies (the fraction of runs assigning two cells to the same inferred clone bin) and consensus memory scores averaged across runs. Row-normalizing the co-clustering matrix gives the pseudoclone probability distribution for each seed cell. (3) Pseudoclone embedding constructs the lineage-informed pCORAL manifold from the inferred pseudoclones and consensus memory genes. (4) Metaclone coarse-graining and dynamic inference reconstruct a lineage-resolved cell-fate map.

Building on previous reports^31,33^ and our own CORAL analysis of lineage-traced data^32^, we propose three principles to guide memory-feature selection (**Figure 1B**): (i) memory genes tend to show high expression or high variance; (ii) co-expressed memory genes can be evaluated as modules; and (iii) bins enriched for clonally related cells can reveal additional memory genes through expression differences across bins.

Driven by these principles, we designed a memory-feature extraction strategy (**Figure 1C**; **Methods**). First, a piecewise mean–variance function identifies a candidate memory-gene set while discarding known confounders such as cell-cycle genes. Second, non-negative matrix factorization (NMF) groups the candidates into putative modules. In the subsequent validation step, the top genes of each module define initial memory bins, and a module is retained only if genes in the broader candidate-gene set display high variance across these bins, as expected from principle (iii) when the module captures genuine clonal structure. Top genes from the retained modules form the key-feature gene set used for pseudoclone inference.

isCORAL represents each cell by a probability distribution over its inferred clonal relatives, termed a pseudoclone (**Figure 1C**). Across randomized runs, the method selects memory features, partitions cells by spectral clustering and records how frequently each cell pair is grouped together. Aggregating these runs yields consensus pseudoclone distributions and gene memory scores. This probabilistic representation accommodates clones spanning multiple cell states without requiring experimental lineage labels.

isCORAL then uses these pseudoclones to construct a clone-oriented embedding space, so that clustering and trajectory inference are performed at the pseudoclone level, yielding cell-fate maps with lineage resolution (**Figure 1C**; **Supplementary Table 1**).

### isCORAL detects memory genes with high accuracy

Memory genes retain expression differences associated with clonal ancestry^30–32^: their expression is more similar within a clone than between clones (**Figure 1B, 2A**). Functionally, these genes retain substantial lineage information and are enriched for drivers of processes such as fate priming^40^ and drug resistance^41^. However, they are not necessarily the most highly variable genes overall, nor do they always separate into distinct transcriptomic clusters. Consequently, they are easily overlooked during classical data exploration and typically require experimental lineage tracing to be uncovered^32^. We first asked whether isCORAL can detect these genes from transcriptomes alone.

Lineage-traced datasets make this evaluation possible: isCORAL infers memory genes from transcriptomes alone, while the experimentally measured lineages supply ground truth. Ground-truth memory strength was quantified as the ANOVA omega-squared statistic (the relative variance among clones), as formalized in our previous CORAL framework^32^. We benchmarked isCORAL’s memory-gene detection (**Figure 2B**) on multiple lineage-tracing datasets spanning developmental models^37,42^ such as hematopoietic stem cell (HSC) differentiation^37^ and cancer models across various timescales^32,43,44^ (**Supplementary Table 2**), with the aim of extending such analyses to barcode-free data, including large-scale atlases and primary tissues. The baselines (**Figure 2C**) comprised published in silico clonal- and memory-inference tools (GEMLI^33^ and an adapted, faster implementation of Power-Seek^36^ (**Methods**)), simplified pseudoclone baselines (SNN clustering with or without cell-cycle regression), and standard feature-selection methods (scran^45^, VST and MVP in Seurat).

**Figure 2.**
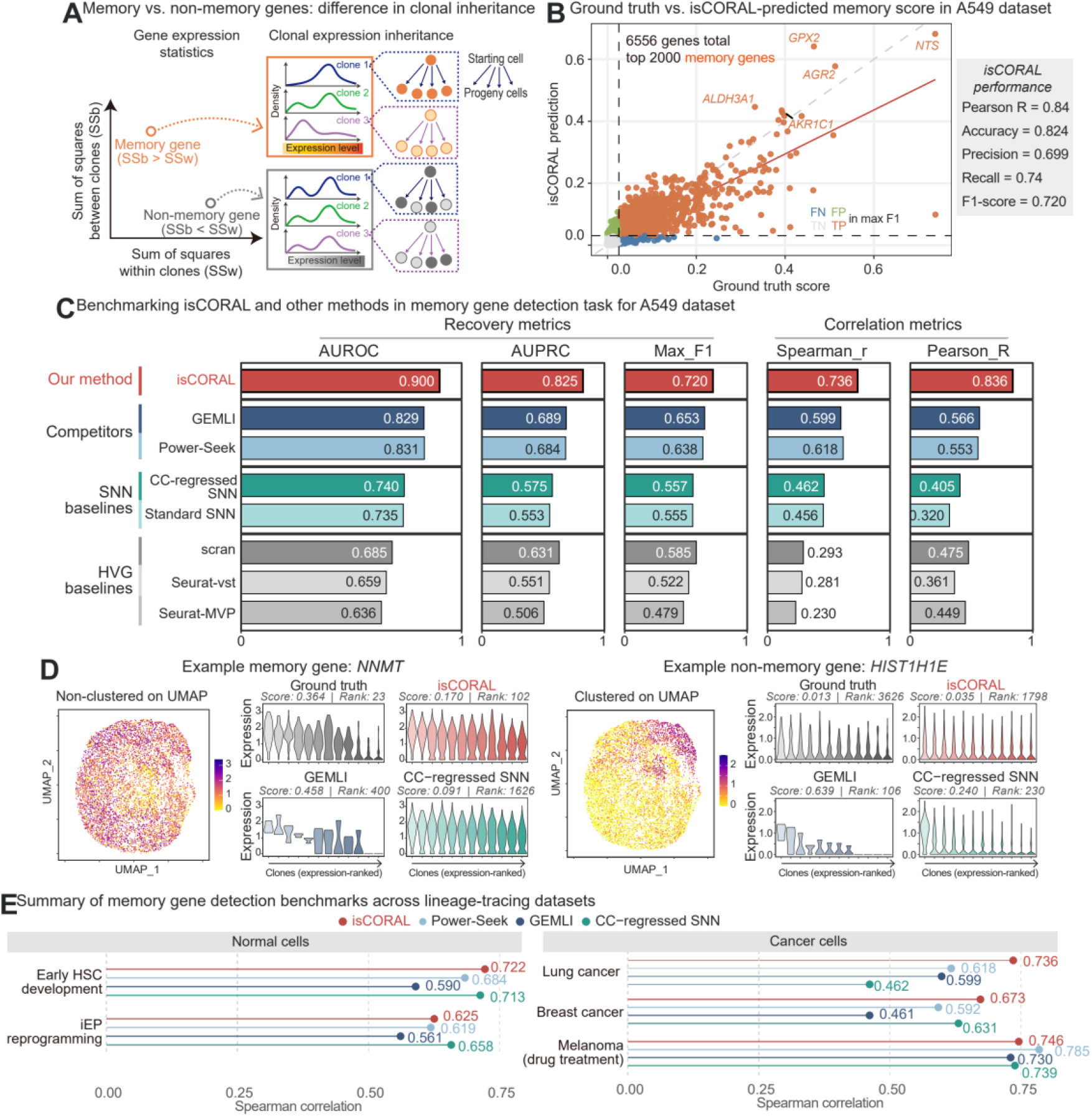
isCORAL detects memory genes with high accuracy. (A) Concept of memory genes in the lineage-tracing framework. When expression differences persist across cell generations, a memory gene varies more between clones than within clones. This property is quantified by the omega-squared statistic, the proportion of variance explained by clonal identity, computed from the between- and within-clone sums of squares (SSb and SSw) in an ANOVA framework; omega-squared serves as the ground-truth score for benchmarking memory-gene detection. (B) Scatter plot of true versus isCORAL-inferred omega-squared values in the A549 lineage-tracing dataset^32^ (8,972 cells; 6,556 genes). Most top-ranked memory genes (top 2,000) are successfully recovered, and the quantification of omega-squared is accurate (Pearson *r* = 0.84). The decision threshold for isCORAL is set to maximize the F1 score. Several top-detected memory genes are highlighted. Orange, true positives; green, false positives; blue, false negatives; grey, true negatives, with positives defined as the top 2,000 ground-truth memory genes. (C) Comprehensive benchmark of memory-gene detection in the A549 lineage-tracing dataset, evaluating GEMLI, a PCA-based approximation of Power-Seek (Methods), shared nearest-neighbor (SNN) baselines (cell-cycle-regressed SNN and standard SNN), and highly variable gene (HVG) baselines (scran, Seurat Mean-Var-Plot (MVP), and Seurat Variance-Stabilizing Transformation (VST)). Recovery metrics (areas under the receiver operating characteristic and precision–recall curves, AUROC and AUPRC, and maximum F1 score for the top 2,000 memory genes) and quantification metrics (Pearson and Spearman correlations) are reported. isCORAL scores highest on all five metrics in this dataset. (D) Case studies of memory-gene detection across representative methods. Violin plots show expression distributions within experimentally recorded or inferred clones, ranked by expression. *NNMT* is a high-memory gene (score = 0.364, rank = 23) despite its dispersed distribution in the UMAP embedding. Both isCORAL (score = 0.170, rank = 102) and GEMLI (score = 0.458, rank = 400) successfully identify it as a high-memory gene, whereas the cell-cycle-regressed SNN substantially underestimates its memory strength (score = 0.091, rank = 1,626) despite the dispersed clonal expression pattern. Conversely, the cell-cycle-associated gene *HIST1H1E* has negligible memory (score = 0.013, rank = 3,626) despite exhibiting clustered expression on the UMAP plot. GEMLI (score = 0.639, rank = 106) and the cell-cycle-regressed SNN (score = 0.240, rank = 230) assign it much higher memory scores, whereas isCORAL (score = 0.035, rank = 1,798) only slightly overestimates its score. (E) Memory-gene detection across lineage-tracing datasets spanning hematopoiesis, reprogramming and cancer. The Spearman correlation between estimated and experimentally measured memory strength is shown for each method and dataset; values are annotated beside the points. Power-Seek denotes the PCA-based approximation described in Methods.

Under standard parameter settings, isCORAL achieved the best overall performance in recovering top memory genes and estimating memory strength (Pearson and Spearman correlations; **Figure 2C–E, Figures S1–S2**). The largest gains were observed in the A549 lung cancer and SUM159 breast cancer datasets, while performance in the remaining datasets was similar to that of the strongest alternatives. isCORAL also recapitulated clone-level gene expression distributions across systems, including dispersed memory patterns that were poorly captured by conventional clustering.

Two case studies in the A549 dataset illustrate how isCORAL distinguishes heritable memory from transient variability. The memory gene *NNMT* is highly scattered across transcriptomic space and is therefore overlooked by classical clustering (**Figure 2D, Figure S1A-B**), whereas isCORAL ranked it closest to its ground-truth memory rank among all methods (rank 102 versus true rank 23) and reconstructed its approximate clonal distribution. Conversely, the highly variable cell-cycle gene^46^ *HIST1H1E* was promoted as a high-memory gene by several baselines even after cell-cycle regression, whereas isCORAL assigned it low memory (**Figure 2D**). Thus, isCORAL recovered inherited expression differences that global transcriptome clustering obscured while assigning low memory to a highly variable cell-cycle gene. The same pattern held in the HSC dataset^37^, where isCORAL successfully distinguished the dispersed memory gene *Pde4d* from the proliferation-associated gene *Top2a* (**Figure S1C-H**), and in the SUM159 breast-cancer^44^, drug-resistant melanoma^43^ and iEP (induced endoderm progenitor) reprogramming^42^ benchmarks (**Figure S2**). In the melanoma dataset, isCORAL detected memory genes whose strength may be underestimated in the experimental ground truth: genes expressed predominantly in small clones, such as *MLANA*, that are sparsely sampled by experimental lineage tracing (**Figure S2G**).

We also introduced SNN-based baselines for memory inference by treating conventional transcriptomic clusters as proxy clones (**Figure 2E**). With cell-cycle regression, this simple approach outperformed the tested specialized methods other than isCORAL in most datasets. It approximates memory when clonal relatives group together by overall expression, but underestimates inherited differences when distinct clones are mixed within clusters, as illustrated by *NNMT*. Its performance depends on preprocessing: without cell-cycle regression, the SNN baseline can fall below GEMLI, which lacks built-in cell-cycle adjustment.

Having established detection accuracy across datasets, we next asked how sensitive isCORAL is to data quality. In downsampling experiments (**Figure S2J**), isCORAL maintained its accuracy even at low cell numbers or UMI depths and still exceeded the full-data performance of the alternative methods. This robustness supports extending isCORAL to the sparsely sampled settings that motivated this benchmark: barcode-free atlases and spatial transcriptomics.

### Pseudoclones recover clonal relationships across changing cell states

After validating memory gene inference, we next tested whether isCORAL could reconstruct clonal relationships from a single transcriptomic snapshot. Despite efforts such as GEMLI^33^, recovering clonal relatives across different cell states remains difficult. The central challenge is that global transcriptomic similarity does not reliably reflect ancestry: unrelated cells can converge on similar states, while clonal relatives can become transcriptionally dissimilar as their states change.

We evaluated reconstruction at the level of clonal distributions: for each seed cell, the inferred pseudoclone should approximate the transcriptomic distribution of its experimentally recorded clonal relatives (**Figure 3A**). To quantify this approximation, we used three complementary classes of metrics (**Figure 3B**): (1) absolute pairwise accuracy (AUROC and AUPRC over lineage pairs); (2) relative accuracy at the distribution level (energy distance between inferred pseudoclones and true lineages); and (3) relative accuracy at the pseudobulk level (L1 distance between inferred and true pseudobulk profiles).

**Figure 3.**
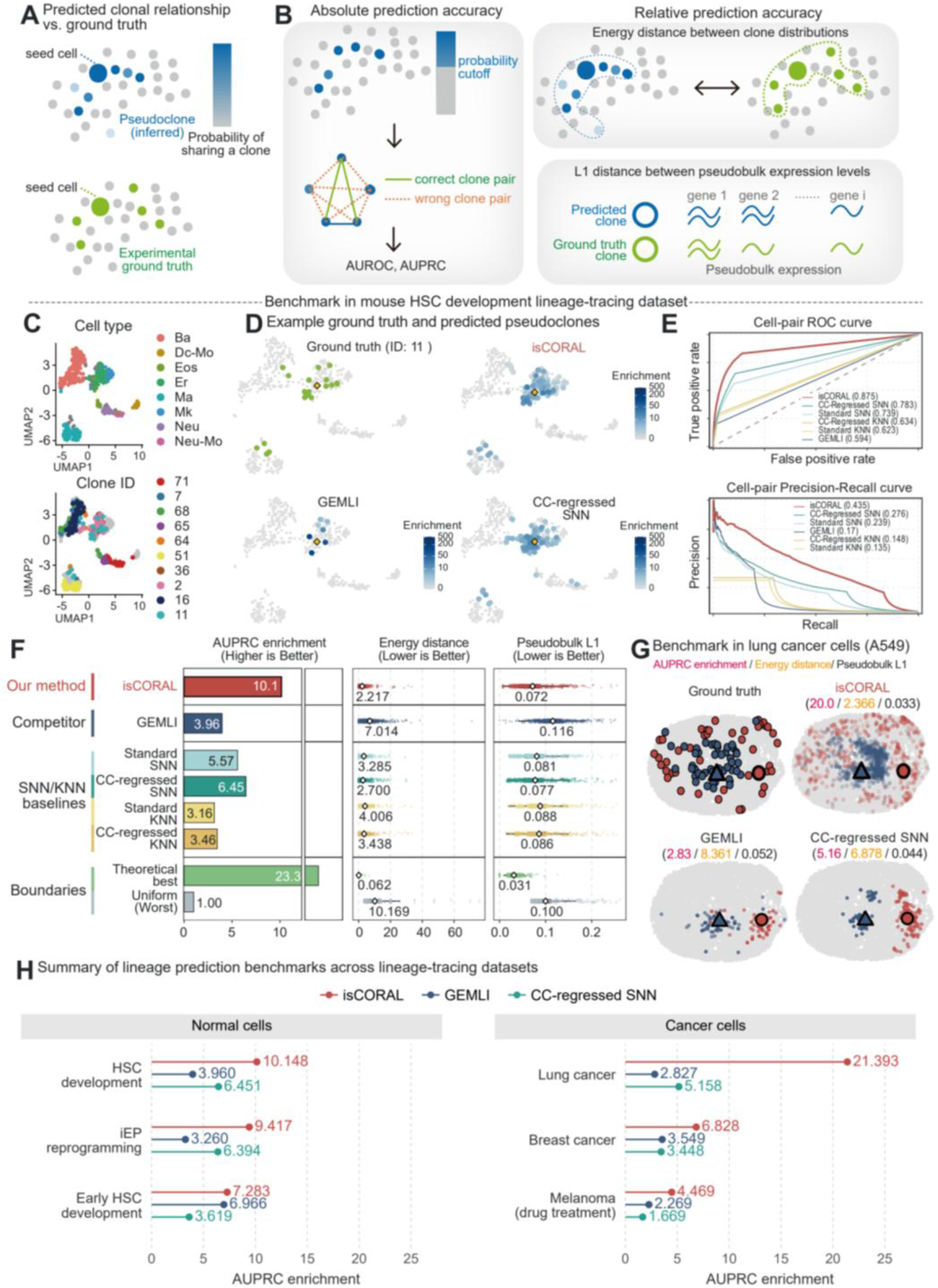
Pseudoclones match experimental lineage-tracing ground truth. (A) Probabilistic pseudoclone reconstruction. For each seed cell, normalized cell-pair co-clustering frequencies assign weights to potential clonal relatives, yielding a pseudoclone distribution (blue). The inferred distribution is compared with the experimentally recorded clone (green). (B) Three measures of reconstruction accuracy. Cell-pair accuracy is evaluated by receiver operating characteristic (ROC) and precision–recall (PR) curves, treating pairs with the same experimental barcode as positives. Distribution accuracy is measured by the energy distance between inferred and recorded clones in the top 30 principal components. Pseudobulk accuracy is measured by the L1 distance between their averaged expression profiles. (C) UMAP visualization of the HSC-differentiation lineage-tracing dataset^28^ (GSE262580, 549 cells) used for benchmarking. Cells are highlighted by their cell types (top) or by their experimental clone barcodes (showing only the 10 largest clones). Cells within a single clone tend to cluster in the transcriptomic space, supporting the feasibility of transcriptome-based lineage inference. Nevertheless, some clones span multiple cell types. (D) Examples of probabilistic pseudoclone estimation (blue) using different methods on a UMAP plot, compared with the ground truth clone from (C). The seed cell is highlighted by a diamond shape. Ground-truth clone 11 is relatively dispersed, increasing the difficulty of lineage inference. isCORAL reconstructs such dispersed, cross-cell-type clones with high accuracy. GEMLI assigns high weights to a smaller subset of clonal relatives in these examples. The new baseline we propose, the SNN graph with cell-cycle regression, also reconstructs clones reasonably well, although its inferred distributions include more cells outside the recorded clone. (E) ROC and PR curves of competing pseudoclone inference methods for transcriptome-based clone inference, using the dataset from (C). isCORAL consistently outperforms all baselines, with its performance curves nearly enveloping the others in both ROC and PR spaces. GEMLI displays high precision at low recall but fails to outperform the SNN baselines at higher recall. The nearest-neighbor (KNN) baselines have the lowest areas under the curves in this dataset. (F) Reconstruction performance measured by AUPRC enrichment over uniform allocation, energy distance and pseudobulk L1 distance. Higher AUPRC enrichment and lower distances indicate better recovery. Each point in the distance plots represents one seed-cell pseudoclone compared with its recorded clone. Uniform allocation and the theoretical-best references are defined in Methods. isCORAL has the highest AUPRC enrichment and the lowest energy and pseudobulk distances among the tested methods in this dataset. (G) Representative pseudoclone reconstruction in the lung cancer cell line A549^32^ (8,972 cells), comparing experimental clones with isCORAL, GEMLI and cell-cycle-regressed SNN. The outlined red circle and blue triangle mark the seed cells. Dot colors distinguish the two lineages; inferred pseudoclone enrichment is shown relative to uniform allocation (Methods). The three annotated scores are AUPRC enrichment, energy distance and pseudobulk L1 distance, respectively. (H) Summary of pseudoclone reconstruction across datasets spanning hematopoiesis, reprogramming and cancer, showing AUPRC enrichment for the indicated methods. isCORAL achieves the highest AUPRC enrichment in the displayed datasets. Additional early-HSC examples^37^ are shown in Figure S3.

We next benchmarked pseudoclone construction methods on a public single-cell RNA-seq dataset of mouse HSC development with ground-truth lineages^28^ (**Figure 3C**). isCORAL was compared with GEMLI and standard Seurat SNN/KNN baselines; for these graph-based methods, pseudoclones were generated from their cell–cell graphs. isCORAL substantially outperformed all baselines across both absolute and relative metrics (**Figure 3D-F**). In the dispersed-clone examples, GEMLI assigned high weights to only a small subset of recorded relatives, whereas the SNN baseline included more cells outside the recorded clone (**Figure 3D**). GEMLI showed high precision at low recall but lost precision rapidly as recall increased (**Figure 3E**). isCORAL combined precise local relationships with broader clone recovery, achieving the best pairwise, distribution and pseudobulk accuracy in this benchmark (**Figure 3E,F**).

We then evaluated these methods on larger lineage-tracing datasets spanning different biological contexts (**Figure 3G,H, Figures S3-S5**). These tasks are particularly challenging owing to highly imbalanced positive-to-negative pair ratios; we focused primarily on AUPRC enrichment and relative accuracy. isCORAL performed best across nearly all datasets, with two-to four-fold improvement over the strongest baseline in some cases. The tested contexts spanned hematopoiesis^37^ (**Figure S3**), cancer plasticity^32,44^ (**Figure 3G,H, Figure S4**), drug response^43^ (**Figure S5A-C**) and cellular reprogramming^42^ (**Figure S5D-F**); isCORAL performed robustly in all of them. Existing methods failed particularly in tumor-related tasks, where clones are highly plastic and overlap substantially in global transcriptomic space, whereas isCORAL reconstructed these dispersed clonal distributions without additional parameter tuning.

Together, these benchmarks show that isCORAL recovers clonal expression distributions across distinct cell states more accurately than the tested alternatives. The identified memory genes provide a molecular basis for interpreting these relationships, while the reconstructed distributions enable analysis of fate differences obscured by global transcriptomic similarity.

### Pseudoclone embeddings reveal early fate bias and multilineage clones

We next asked whether inferred clonal relationships could reveal fate differences hidden within conventional cell state maps. Motivated by recent clone-embedding strategies^32,47,48^ developed for lineage-tracing data, we tested whether a pseudoclone-driven embedding could likewise support lineage-tracing-free cell-fate research. We therefore designed a pseudoclone CORAL (pCORAL) strategy as a key analytical module of isCORAL (**Figure 4A**), adapted from our previous clone-embedding algorithm CORAL. pCORAL uses the inferred pseudoclones to generate a joint embedding of pseudoclones and cells. Briefly, we computed a cell-cell energy-distance matrix from the inferred pseudoclones and applied classical multidimensional scaling (MDS), followed by regression onto memory-gene expression, to construct a space that separates cells according to their pseudoclone distributions rather than global transcriptomic similarity (**Methods**).

**Figure 4.**
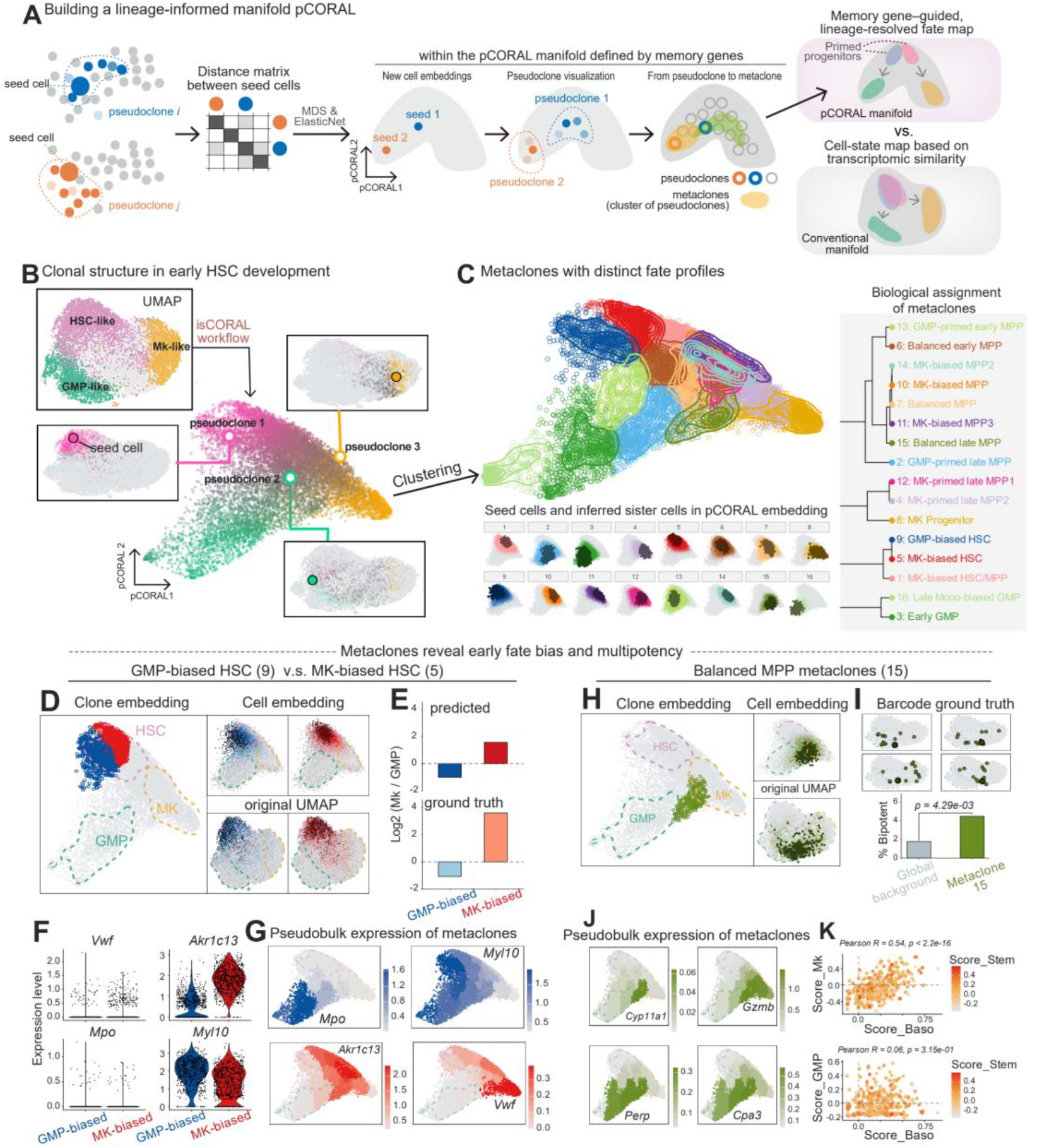
Pseudoclone embeddings reveal early fate bias and multilineage clones. (A) Workflow for building the lineage-informed pCORAL manifold. Distances between seed cells are defined by the squared energy distance between their inferred pseudoclone distributions, computed in the normalized expression space of the top 3,000 inferred memory genes. Multidimensional scaling (MDS) gives initial coordinates, and elastic-net regression predicts the coordinates from memory-gene expression. Probability-weighted averaging then generates pseudoclone coordinates; clustering groups pseudoclones into metaclones. Schematics at right compare a fate map organized by pseudoclone relationships inferred from memory-gene expression with a conventional map organized by cell-state similarity. (B) Clonal structure in an early HSC-development dataset (in vitro, day 7 after sorting). The conventional UMAP shows HSC, granulocyte–macrophage progenitor (GMP) and megakaryocyte (MK) identities after cell-cycle regression (markers in Figure S6A). The pCORAL embedding is reconstructed from isCORAL pseudoclones without experimental lineage labels. Hollow circles represent inferred pseudoclones, colored by their weighted cell-type proportions. Insets show representative seed cells and their inferred sister-cell distributions in conventional UMAP; paired LARRY validation is shown separately in Figure S6B,D. (C) Visualization of metaclone clustering derived from pseudoclone embeddings. (Top) In the pseudoclone embedding, hollow circles represent individual pseudoclones, colored by their assigned metaclone identity. (Bottom) Faceted cell embedding plots display individual metaclones; seed cells are highlighted in darker colors, while inferred sister fates are shown as lighter, transparent dots. (Right) Metaclone nomenclature and the hierarchical clustering tree are based on the combined distributions of seed cell types and their sister fates (Figure S7). (D) An example of metaclone biological relevance, where metaclone 5 represents MK-biased HSCs (red) and metaclone 9 represents GMP-biased HSCs (blue). The pseudoclones are highlighted in the pCORAL clone embedding, while the seed cells and relevant sisters are highlighted in both the pCORAL cell embedding and the UMAP space. The dashed-line boxes in each embedding indicate the high-density area of specific cell types. Compared to the original UMAP, the pCORAL embedding better separates the two types of clones. (E) The predicted and actual sister cell-fate biases of metaclones 5 and 9, represented as log2[(MK proportion + 1%)/(GMP proportion + 1%)]. According to the LARRY lineage-tracing ground truth, metaclone 5 is biased toward the MK fate and metaclone 9 toward the GMP fate; the inferred and recorded sister-cell compositions show the same direction of fate bias. (F) Violin plots showing classical MK/GMP markers and the seed cell markers for metaclones 5 and 9. Typical MK/GMP markers are rarely expressed in either metaclone, consistent with their HSC annotation. In contrast, several other markers are differentially expressed in the seed cells of the two metaclones. (G) Metaclone-level pseudobulk expression of the genes in (F), projected onto the pCORAL clone embedding. The classical MK/GMP markers and early HSC fate-bias markers show distinct distributions. (H) A second example of metaclone biological relevance: metaclone 15, classified as balanced late multipotent progenitors (MPPs), has inferred sister fates spanning both the MK and GMP compartments. The pseudoclones are highlighted in the pCORAL clone embedding, while the seed cells and relevant sisters are highlighted in the pCORAL cell embedding and UMAP space. The seed cells and inferred sisters are dispersed in conventional UMAP but connected by their pseudoclone relationships. (I) Validation of the inference in (H) using the LARRY lineage-tracing ground truth. The four largest relevant LARRY clones (top) are plotted; their cells span both GMP and MK compartments. The bottom plot compares the proportion of cells belonging to recorded clones containing both GMP and MK cells in metaclone 15 versus all other metaclones (two-sided Fisher’s exact test, P = 4.29 × 10^−3^). (J) Metaclone-level pseudobulk expression of representative genes differentially expressed between metaclone 15 and all other metaclones, projected onto the pCORAL clone embedding. (K) Single-cell gene-set scores in metaclone 15: basophil/mast-cell score versus MK score (top) or GMP score (bottom). Each dot represents a seed cell; color indicates the HSC stemness score. Pearson correlations and two-sided P values are annotated.

Specifically, we applied isCORAL to the day-7 (D7) subset of an in vitro HSC-expansion single-cell RNA-seq study^37^ and asked whether the pseudoclone embedding resolves fate structure that conventional analyses leave hidden (**Figure 4B**). Although the original study included LARRY-based lineage barcodes, we used them only for validation. Classical embedding strategies, such as PCA followed by UMAP, generally reflect key cell states during hematopoiesis, including the HSC, megakaryocyte (MK), and granulocyte–macrophage progenitor (GMP) states (**Figure S6A**). However, the transition paths between these states remained unresolved, limiting trajectory-level inference. isCORAL instead used the inferred pseudoclone distribution of each seed cell to generate pCORAL embeddings of pseudoclones and cells (**Figure 4B, Figure S6B-D**). In this embedding, undifferentiated clones, fate-restricted clones, and multi-fate clones collectively form a cell-fate map at lineage resolution.

To systematically summarize these pseudoclones, we performed Leiden clustering of the pseudoclone embedding to extract recurring clone states, which we termed “metaclones” (**Figure 4C**). In the Waddington-landscape metaphor, the region where the cells of a clone tend to localize corresponds to a putative attractor, or valley. The cell embedding allowed us to characterize both seed cells and sister cells at the metaclone level. We defined 16 metaclones during early HSC development, named according to their seed-cell and inferred sister-cell distributions (**Figure 4C**). This metaclone structure was consistent with both the LARRY ground truth (**Figure S6E-F**) and the known hematopoietic hierarchy^49^ (**Figure S7A-D**), suggesting that the embedding and clustering steps further stabilize the downstream inference.

Importantly, metaclone analysis revealed fate bias without any experimental lineage-tracing information. For example, we identified two types of HSCs with distinct fate preferences despite highly overlapping UMAP distributions (**Figure 4D**). The GMP bias of metaclone 9 was accurately predicted by isCORAL, whereas metaclone 5 showed an even stronger MK preference than predicted, potentially reflecting the higher proliferation rates of MK-biased fates (**Figure 4E**). Next, we performed differential gene-expression analysis (**Figure 4F, Figure S7E**) on seed cells from these two metaclones to identify markers that distinguish these fate-biased HSC populations. Because both metaclones 5 and 9 consist primarily of HSCs, they rarely express typical GMP or MK markers (for example, *Vwf* and *Mpo*). In contrast, we identified several early fate-bias markers, including *Akr1c13*, which is known to be highly expressed in basophil/mast populations but not typically in MKs^50^. We found *Akr1c13* to be enriched in HSC and multipotent progenitor (MPP) states with MK-biased fates, though expression was lost in mature MK progenitors (**Figure 4G**). Thus, isCORAL revealed early cell-priming events without experimental lineage labels as input.

Beyond fate priming, metaclone analysis also resolves multipotent lineages. As an example, we focused on metaclone 15, which displayed a low HSC proportion but little preference for GMP or MK fates (**Figure 4H**). The inferred sister cells are highly scattered across multiple cell types in the original UMAP; the prediction that these cells are clonally related is therefore not evident from global transcriptomic similarity. Yet this prediction was supported by the corresponding experimental ground-truth clones (**Figure 4I**), which displayed a scattered distribution and significant GMP/MK bipotent fates. Further differential-expression analysis (**Figure 4J, Figure S7F**) highlighted the complexity of these cells, identifying genes associated with distinct processes: *Cyp11a1* in lipid metabolism^51^, *Perp* as a TP53 apoptosis effector^52^, and *Gzmb* and *Cpa3* as indicators of early basophil/mast cell fate^53^. Some of these genes were also highly expressed along the GMP-biased trajectory, and others along the MK-biased trajectory, suggesting that metaclone 15 may sit at the crossroads of the GMP/MK fate decision, close to the basophil/mast fate. Single-cell expression analysis showed that MK and basophil/mast gene programs were positively correlated within metaclone 15 (**Figure 4K**), supporting their coexistence within the same transcriptional state. This aligns with recent reports that the basophil/mast fate can be coupled with MK fates rather than with the classical GMP fate of standard hematopoietic models^54–56^.

Together, these analyses distinguish fate-biased HSC populations from clones spanning multiple developmental compartments, relationships that conventional transcriptomic embeddings obscure.

### Metaclone transitions predict HSC developmental routes and fate probabilities

We next asked how the clonal relationships resolved by isCORAL could predict developmental routes and the probabilities of alternative fates. We therefore constructed a transition model linking metaclones.

We observed that metaclone structures naturally delineate coarse-grained developmental trajectories (**Figure 4C**): intuitively, cells are more likely to differentiate into, or originate from, the states in which their sister cells reside. We therefore modeled transitions between metaclones (**Figure 5A**). Multiple pseudotime metrics could be applied in this framework to establish directionality; here, we leveraged the CytoTRACE2 score^57^, derived solely from single-cell RNA-seq data, as a measure of developmental potential. Although single-cell CytoTRACE2 scores are noisy (**Figure S8A**), metaclone-level scores aligned well with the metaclone biology described above (**Figure 5B**). We thus used these scores to constrain transition directions, allowing flow only from high-to low-potential metaclones (**Figure S8B**). Overlap between one metaclone’s inferred sister cells and another’s seed cells defined transition strength, while decreasing developmental potential defined direction. These constraints yielded a directed acyclic graph (DAG) of predicted developmental transitions.

**Figure 5.**
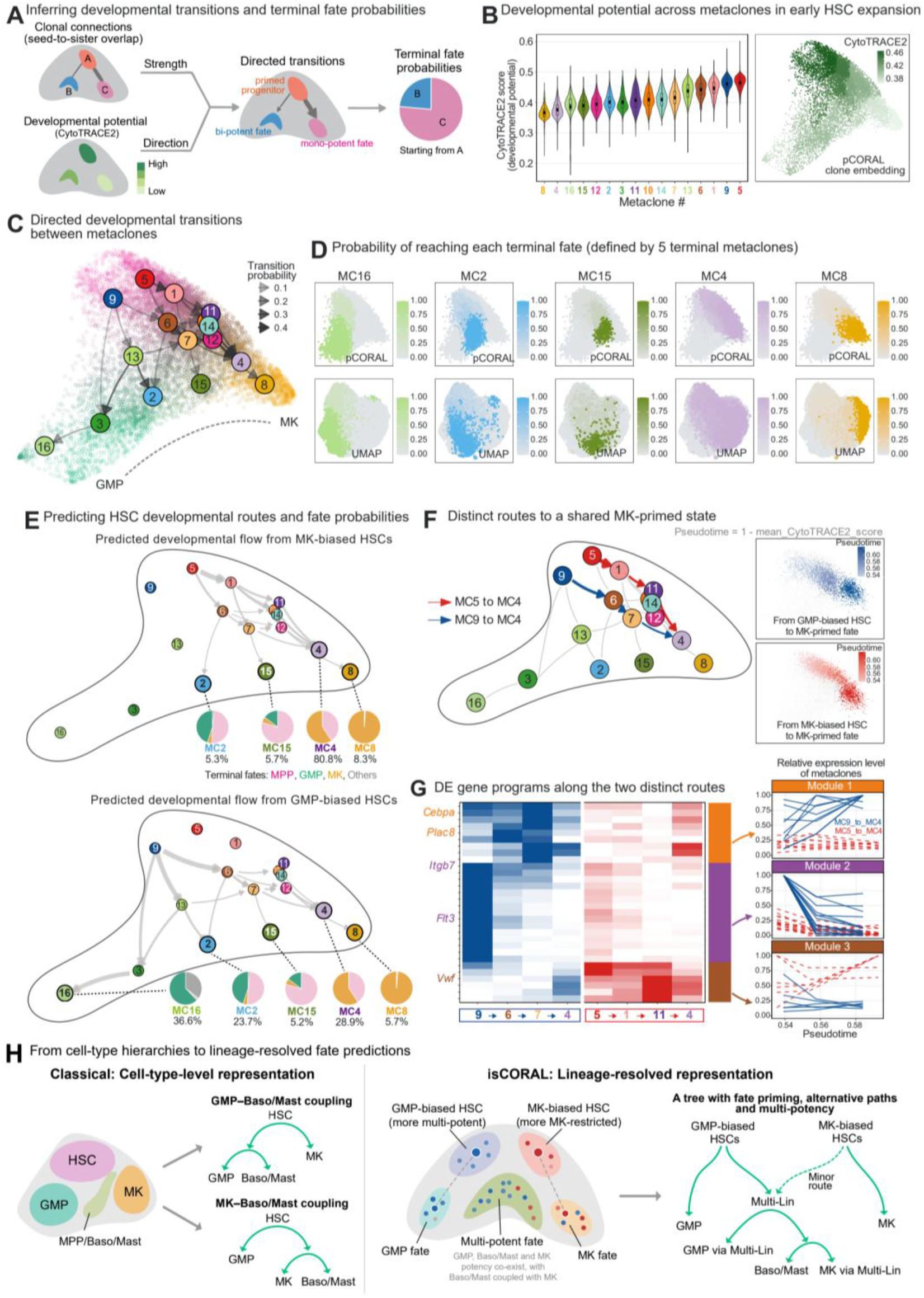
Metaclone transitions predict HSC developmental routes and fate probabilities. (A) Workflow for inferring metaclone transitions and terminal-fate probabilities. By integrating the pseudoclone-derived metaclone structure with developmental potential estimation (CytoTRACE2), isCORAL infers a directed acyclic graph (DAG) of metaclones. Edge weights are estimated from the overlap between one metaclone’s sister cells and another’s ‘seed’ cells, while edge directions are determined by the drop in developmental potential. Filtering and normalization yield directed transition probabilities; these are used to compute the probability of reaching each terminal metaclone. (B) Developmental potential estimated by CytoTRACE2 in the early HSC-expansion dataset. Left, seed-cell score distributions in each metaclone, ordered from low to high mean score. Right, mean seed-cell scores for each metaclone projected onto the pCORAL clone embedding. Higher scores indicate greater predicted developmental potential. (C) Filtered directed transition graph of early HSC development. Background pseudoclone dots are plotted as in Figure 4B, reflecting the cell-type distribution in the pCORAL clone embedding; numbered circles identify metaclones. Arrows indicate directed transitions; transparency reflects transition probabilities. (D) Predicted probabilities of reaching the five terminal metaclones (16, 2, 15, 4 and 8) identified in (C). Each probability is conditional on the starting metaclone and can be nonzero for multiple reachable terminal fates. The top row displays these probabilities in the pCORAL embedding; the bottom row shows the same probabilities in conventional UMAP. Terminal metaclones have no outgoing transitions after the graph-filtering step. (E) Predicted transition paths originating from MK-biased HSCs (metaclone 5) and GMP-biased HSCs (metaclone 9) in the pCORAL embedding. Arrows denote transition trajectories, with line widths proportional to the expected transition flux. For each reachable terminal metaclone, the percentage gives the absorption probability from the indicated starting metaclone, and the pie chart shows its seed-core cell-type composition (pink, MPP; green, GMP; orange, MK; gray, others). (F) Comparison of the two most probable converging transition paths from HSCs to the shared MK-primed state: metaclone 5-1-11-4 (red, from MK-biased HSCs to the MK-primed state) and metaclone 9-6-7-4 (blue, from GMP-biased HSCs to the MK-primed state), shown in the DAG (left) and the pCORAL clone embedding (right). In the embedding space, colors represent pseudotime defined as one minus the mean seed-cell CytoTRACE2 score of each metaclone. (G) Dynamic expression patterns of differentially expressed genes along paths 5-1-11-4 (red) and 9-6-7-4 (blue). The top 30 representative genes showing major expression divergence along these paths are clustered into three distinct modules, visualized via heatmap (left) and individual gene-expression trajectories (right; blue solid lines, 9-to-4 path; red dashed lines, 5-to-4 path). Expression is averaged across seed cells within each metaclone; each gene is normalized to its peak value across both paths. Module 1 shows persistent residual GMP-relevant memory along the ‘blue’ path. Module 2 shows GMP-relevant priming signals that disappear early along the ‘blue’ path. Module 3 shows MK-relevant priming signals that appear earlier along the ‘red’ path. (H) Conceptual comparison of cell-type-level developmental representations (left) with the lineage-resolved isCORAL representation (right). Metaclones distinguish GMP-biased and MK-biased HSCs, resolve fate priming and alternative routes through a multilineage progenitor state, and represent clones with multiple fates. The schematic summarizes these lineage-specific predictions rather than proposing a new hierarchy of cell types. Multi-Lin denotes an inferred progenitor state with multiple predicted lineage outputs. The dashed connection denotes the minor predicted route.

After unsupervised filtering of weak transitions (**Figure S8B**), we obtained a directed map of early HSC development (**Figure 5C**), which provides a quantitative picture of how HSCs determine their fates within a discrete, hierarchical, yet probabilistic model. For example, we defined five terminal metaclones that either primarily contain a single fate (metaclone 16 is predominantly GMP) or encompass multiple fates (metaclones 2 and 15 contain both MK and GMP fates); the probabilities of reaching each terminal fate can then be estimated (**Figure 5D**).

The dynamic fate map adds a temporal dimension to pseudoclone inference. Because transcriptional memory typically persists over short timescales, pseudoclones themselves capture recent clonal relationships; the map instead chains directed transitions between metaclones, yielding quantitative trajectories of longer-term fate choice. The two metaclones discussed above, metaclone 5 (MK-biased) and metaclone 9 (GMP-biased), display distinct paths in the dynamic fate map (**Figure 5E**). isCORAL predicted that MK-biased HSCs predominantly reach MK-like terminal fates (e.g., metaclone 4 or 8), with a lower probability of reaching bipotent MK-GMP fates (e.g., metaclones 2 and 15). In other words, although an MK-biased HSC clone may produce GMPs, it is unlikely to become a clone consisting exclusively of GMP cells. By contrast, GMP-biased HSCs were predicted to reach all defined terminal fates, but show a strong preference for GMP-containing metaclones such as metaclones 2 and 16. These inferences are consistent with clonal analyses showing that platelet-biased and monopotent MK-restricted progenitors arise directly from HSCs^58^, with the heritable lineage imprinting reported in barcoded hematopoietic cultures^17,37^, and with “MultiLin” cell states within the classical common myeloid progenitor CMP/GMP gate^59^.

We next asked whether distinct early fate biases could lead to a shared developmental state through different molecular programs. We compared the two dominant trajectories entering the MK-primed fate (metaclone 4): the metaclone 5-to-4 path (5-1-11-4) and the metaclone 9-to-4 path (9-6-7-4). Although these trajectories overlap substantially in UMAP space (**Figure S8C**), they form two parallel paths that are clearly distinguishable in the pCORAL embedding and the DAG (**Figure 5F**). Certain genes were differentially expressed along the two paths and displayed three main patterns, despite both paths leading to similar fates (**Figure 5G, Figure S8D**). Genes in the first group, including the master myeloid regulator *Cebpa*, remained expressed along the 9-to-4 path, indicating strong residual myeloid/GMP memory. Genes in the second group, including the GMP-biased HSC markers *Itgb7* and *Flt3*, were rapidly downregulated during MK differentiation. Genes in the third group, primarily MK-fate markers such as *Vwf*, were upregulated much earlier along the 5-to-4 path than along the 9-to-4 path. These dynamics mirror established regulatory logic: *Vwf* marks platelet-primed HSCs at the apex of the native hierarchy^58^, *Flt3* upregulation accompanies the loss of self-renewal at the stem-cell-to-progenitor transition^60^, and *Cebpa* drives the granulocyte–monocyte program in myeloid progenitors^61^. These inferred trajectories suggest that the timing of gene regulation during hematopoietic fate determination can be lineage-specific. Such path-specific timing is consistent with the early cross-antagonism between megakaryocyte–erythroid and granulocyte– monocyte programs described in studies of hematopoietic commitment^37^. Thus, while these paths overlap in the conventional transcriptomic embedding, isCORAL resolves distinct routes to a shared MK-primed state and identifies the gene expression programs that distinguish those routes.

Overall, isCORAL reconstructs a lineage-aware, dynamic developmental map solely from scRNA-seq data, explicitly modeling early stem-cell priming, alternative developmental paths, and multipotent clones (**Figure 5H**). The map retains clonal relationships across cell states. For example, whether the basophil/mast fate of HSCs is coupled to the MK fate or the GMP fate has long been debated (**Figure 5H, Figure S8E**). The classical hematopoiesis model places basophils and mast cells downstream of GMPs, whereas single-cell studies have observed transcriptomic and clonal-fate coupling between basophil/mast and MK/erythroid fates^17^. Different computational tools have reported evidence supporting both models, even when using the same lineage-tracing ground-truth data^62,63^. The isCORAL map offers a view that reconciles both models: the basophil/mast fate, positioned downstream of a multilineage progenitor state (Multi-Lin; metaclones 15 and 2), is mutually exclusive with the early-determined MK fate (and therefore GMP-coupled at early stages) but strongly coupled with MK cells derived from MPPs. This accounts for the evidence supporting both the MK-coupled and GMP-coupled models, thereby generating specific predictions for future experimental validation. Through this end-to-end lineage-aware modeling strategy, isCORAL provides a powerful tool for dissecting complex developmental dynamics.

### isCORAL reconstructs clonal relationships and developmental trajectories from spatial transcriptomes

While spatial transcriptomics enables in situ molecular profiling, adding experimental lineage tracing to spatial measurements remains technically demanding and is currently infeasible in most human primary tissues. We therefore asked whether isCORAL could extract clonal information computationally from spatial transcriptomic data.

First, we applied isCORAL to a public Slide-tags dataset of *Kras*-mutant mouse lung tumors (**Figure 6A**). The data were generated with KPTracer^38^, a Cas9-based lineage-tracing system that records a lineage tree. Our goal was to test whether the isCORAL pseudoclones faithfully recovered the clonal expression structure recorded in this tree. Specifically, we matched each pseudoclone to its most similar branch in the experimental tree and calculated their transcriptomic energy distance (**Figure 6B**). Smaller energy distances indicate better reconstruction of the clonal expression distribution. isCORAL recovered clonal structure more faithfully than both a random pseudoclone control (median energy distance, 1.42 versus 5.14) and the cell-cycle-regressed SNN baseline (median, 4.18) (**Figure 6B**). The embedding and metaclone-generation steps also recovered the coarse-grained subclone structure of the Cas9-derived lineage tree (**Figure 6C, Figure S9A, B**), allowing us to analyze cancer-relevant metaclones in their spatial context (**Figure S9C**). Thus, isCORAL recovered clonal expression distributions and coarse subclone organization from spatial transcriptomes, with the recorded lineage tree used only for validation.

**Figure 6.**
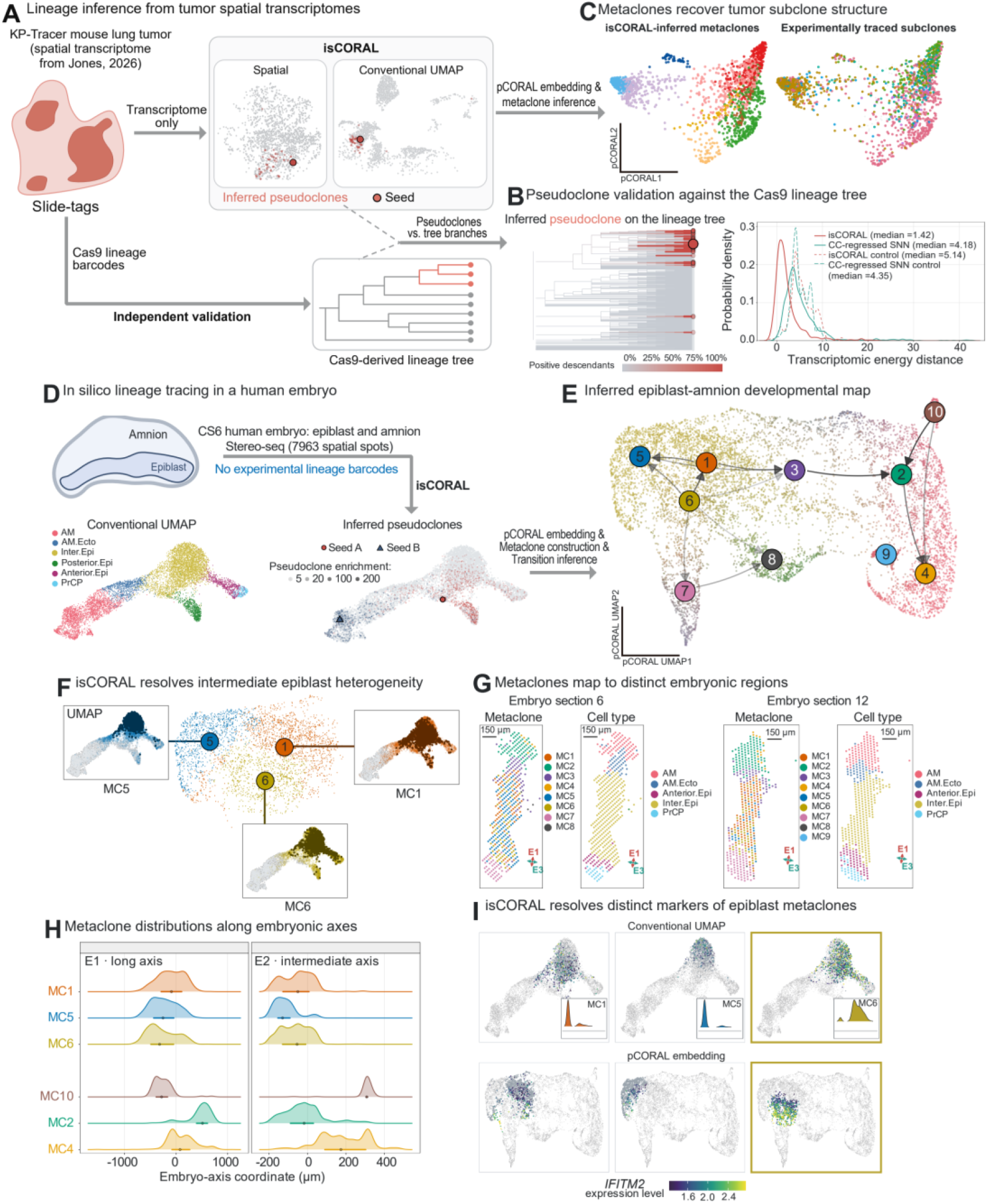
isCORAL reconstructs lineage-resolved fate maps from spatial transcriptomes. (A) Overview of the isCORAL workflow applied to the mouse tumor evolution dataset^38^. The dataset was generated using the Slide-tags technique in a KPTracer lung cancer mouse model. Tumors were driven by oncogenic *Kras/Trp53* mutations, with a Cas9-based lineage-tracing module generating lineage trees. isCORAL reconstructed pseudoclones from the transcriptomic profiles; the recorded Cas9 lineage tree was used for validation. (B) An example of pseudoclones mapped to the Cas9-based lineage tree (left) and quantification of reconstruction accuracy (right), measured by the transcriptomic energy distance between pseudoclones and branches selected to maximize the F1 score of pseudoclone recovery, compared with a random pseudoclone control and a cell-cycle-regressed SNN baseline. Note that the random pseudoclone control was calculated separately for isCORAL and SNN, as their paired branches can differ. Energy distances are computed in the top 30 transcriptomic principal components. (C) Comparison between isCORAL-derived metaclones (colored by metaclone identity) and Cas9 lineage-tree-derived subclones (colored by subclone identity), visualized in the pCORAL clone embedding (n = 9 metaclones, n = 6 subclones). (D) Overview of the isCORAL workflow applied to a human embryo spatial transcriptomics dataset without experimental barcodes^39^. The dataset encompasses the epiblast and amnion subsets (7,963 spots) of a Carnegie Stage 6 human embryo, with cell types assigned in the original study. Transcriptomic profiles served as input for isCORAL to reconstruct pseudoclones for downstream analysis. (E) Developmental map of the human epiblast and amnion. Inferred pseudoclones are plotted in the pCORAL clone UMAP embedding. Dot colors represent the weighted cell-type proportions of each pseudoclone. Large numbered circles identify metaclones. Arrows indicate directed transitions; transparency reflects transition probabilities in the directed acyclic graph (DAG). (F) pCORAL clone UMAP embedding and conventional UMAP of representative intermediate epiblast metaclones (metaclones 1, 5, and 6). In the pCORAL clone UMAP embedding, circle colors indicate metaclone identities. In the conventional UMAP embedding, dark and light dots highlight ‘seed’ and ‘sister’ cells, respectively. (G) Spatial localization of metaclones in embryo sections 6 (left) and 12 (right), colored by metaclone identity or the original cell-type annotation. In section 6, distal to the amnion, MC5 is central whereas MC1 and MC6 are peripheral; in section 12, MC1 and MC6 occupy the anterior and posterior regions, respectively. E1, E2 and E3 denote the long, intermediate and short geometric axes of the reconstructed three-dimensional embryo, obtained by covariance eigen decomposition after section alignment and stacking (Methods). Arrows show the projections of E1 (red) and E3 (teal) onto each displayed section. Scale bars, 150 µm. (H) Coordinate (micrometer) distributions of representative metaclones along the long (E1) and intermediate (E2) geometric axes of the reconstructed embryo. (I) *IFITM2* expression in intermediate epiblast metaclones 1, 5 and 6, visualized in conventional UMAP (top) and the pCORAL embedding (bottom). Color indicates normalized *IFITM2* expression; insets show its expression distribution in each metaclone. *IFITM2* is enriched in metaclone 6.

Having validated the workflow against an experimentally measured lineage tree, we next asked whether the same “virtual lineage” strategy could recover clonal dynamics in primary human tissue. We thus applied isCORAL to a spatially resolved transcriptomic dataset of a Carnegie Stage 6 human embryo^39^ profiled by Stereo-seq (**Figure 6D**), for which experimental lineage ground truth is unavailable.

Specifically, focusing on the 7,963 spatial spots assigned to the epiblast and amnion, we inferred pseudoclones and performed metaclone-level dynamic analysis, defining 10 metaclones (**Figure S9D,E**). Furthermore, we calculated the CytoTRACE2 score for each metaclone (**Figure S10A,B**) and inferred a putative developmental map for the epiblast/amnion system, visualized on a UMAP embedding of the pCORAL clone space (**Figure 6E**). Because experimental lineage ground truth is unavailable for this dataset, we evaluated both the biological plausibility and spatial distributions of these inferred structures.

Overall, the metaclones were anchored in, but refined, the original annotation: seed spots of each metaclone were dominated by a single annotated cell type, whereas inferred sister spots crossed annotated cell-type boundaries (**Figure S9E**), revealing clonal coupling invisible in the annotation itself. For instance, metaclones 1, 5, and 6 represented intermediate epiblast states with high developmental potential (**Figure 6F**), and isCORAL predicted distinct lineage biases toward amniotic or non-amniotic fates for each. This observation was also supported by the spatial distribution of these metaclones: metaclone 5 was located on the side of the epiblast distal to the amniotic cavity, whereas metaclones 1 and 6 occupied the anterior and posterior epiblast, respectively (**Figure 6G,H**). Thus, epiblast populations that intermix in conventional UMAP occupy distinct regions of the embryo and carry distinct predicted lineage biases.

Furthermore, we identified marker genes for these metaclones as candidate priming markers (**Figure S10C**). For example, IFITM2 (interferon-induced transmembrane protein 2), an antiviral protein that restricts membrane fusion^64^, was highly expressed throughout metaclone 6 (**Figure 6I**) and marks a candidate priming trajectory toward non-amniotic lineages: metaclone 6 captured 79% of IFITM2-high spots (227 of 287), whereas the best cell-cycle-regressed SNN cluster combination captured only 28% (79 of 287), indicating that this primed subpopulation is largely invisible to global transcriptome-based clustering (**Figure S10D**). By contrast, genes upregulated in metaclone 1 were enriched for terms related to muscle cell differentiation, endocrine hormone secretion and regulation of STAT tyrosine phosphorylation (**Figure S10E**).

Similarly, for the amniotic cells, metaclones 2, 4 and 10 delineated three distinct subtypes with divergent CytoTRACE2 scores (**Figure S10A**,**F**). Each metaclone was also localized to a specific area of the amnion (**Figure S10G**). The identification of metaclone 10 was particularly informative: it marks a relatively rare amniotic population with high developmental potential and high expression of *WNT9B* and *IGFBP7* (**Figure S10H**), alongside other genes relevant to mesenchymal differentiation (**Figure S10I**).

Together, these analyses resolve spatially distinct epiblast and amniotic populations and predict their lineage relationships without experimental barcodes. The predicted routes and priming markers nominate specific clonal behaviors for testing by targeted lineage tracing, live imaging or perturbation experiments.

## Discussion

isCORAL makes lineage information accessible in standard transcriptomes by identifying memory genes and estimating probabilistic clonal relationships. Building on studies of transcriptional memory^31,32^ and memory-based lineage prediction^33^, it improves the recovery of clonal expression distributions across diverse developmental and cancer systems. These relationships support lineage-resolved embeddings and quantitative fate mapping without experimental lineage barcodes. Integrating transcriptomic memory with genomic and epigenomic measurements^65^ could further connect lineage reconstruction across timescales.

A second contribution is the representation on which isCORAL builds: a probabilistic cell-cell graph that assigns each cell a distribution over its clonal relatives and can be used directly in standard single-cell analyses. This representation parallels the cell– cell similarity graphs of classical scRNA-seq workflows, yet its topology aligns far better with experimental ground truth than KNN/SNN graphs. Cells with shared ancestry can diverge in expression, whereas unrelated cells in similar states can appear transcriptionally similar^29^. Global expression similarity therefore does not reliably distinguish current cell state from ancestry. Pseudoclone-driven embeddings organize cells according to their inferred clonal relationships, revealing fate biases and multilineage configurations obscured by global expression similarity. Combined with estimates of developmental potential, this representation supports directed models of developmental routes and terminal fate probabilities. Clone-embedding strategies such as Clone2vec^48^, ClonoTrace^47^ and our own CORAL^32^ were developed for lineage-tracing data; here, we extend this strategy to barcode-free single-cell analysis. We anticipate that integrating emerging quantitative dynamics-inference strategies (Neural ODEs^66^, optimal transport^15,67^ etc.) with clone or pseudoclone embeddings will further improve the prediction of lineage-level cell-state transitions, a direction already supported by our results with a proof-of-concept transition model.

Biologically, isCORAL resolves fate heterogeneity within shared cell states and connects this heterogeneity to alternative developmental routes. In HSCs, it separates early MK and GMP biases, identifies clones spanning multiple fates, predicts distinct routes to a shared MK-primed state, and reconciles apparently conflicting models of basophil and mast cell fate coupling. These findings show how inferred clonal relationships can reveal developmental organization that is obscured in conventional cell state maps. The mouse tumor application validates inferred clonal structure against an experimentally recorded lineage tree, while the human embryo application illustrates its use where prospective lineage tracing is impractical. Therefore, the strategy could be extended to other systems with lineage-level dynamics, including cancer evolution^68^, reprogramming in regenerative medicine^42^ and immune cell development. Alongside these applications, isCORAL provides memory gene expression and inferred clonal relationships as potential inputs for virtual cell models^2,3^. These features enable tests of whether incorporating lineage memory improves predictions of cell fate and perturbation responses among cells with similar measured states.

**Figure S1.**
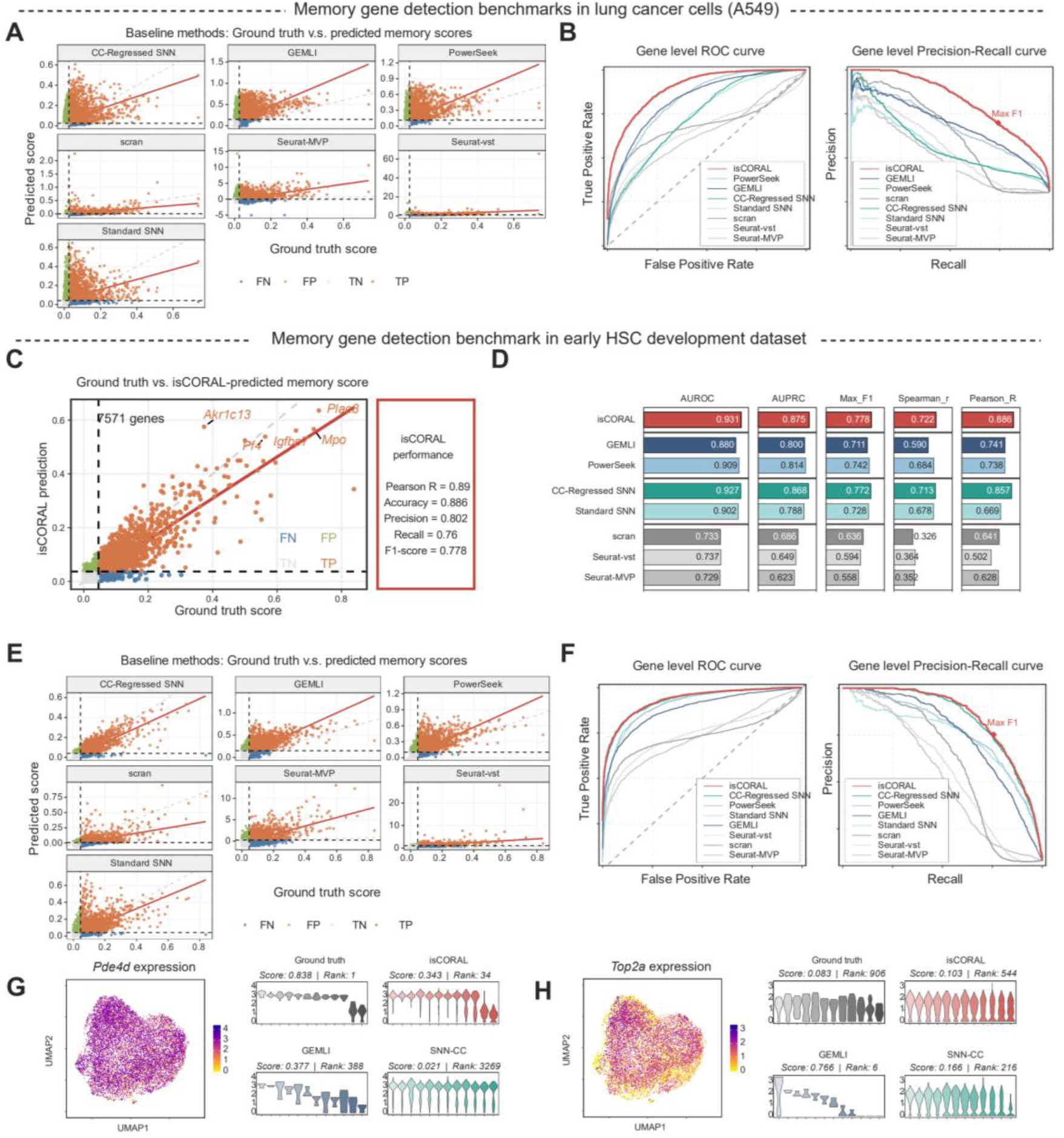
Memory-gene detection benchmarks in the A549 and HSC datasets. (A) Scatter plot of ground-truth versus baseline-inferred memory scores in the A549 lineage-tracing dataset^32^. Dashed lines mark the ground-truth and predicted decision thresholds; orange, true positives; green, false positives; blue, false negatives; grey, true negatives. (B) Receiver operating characteristic (ROC) and precision–recall (PR) curves of different methods for memory-gene detection in the A549 lineage-tracing dataset. The point corresponding to the maximum F1 score of isCORAL is highlighted. (C) Scatter plot of true versus isCORAL-inferred *ω*^2^ (omega-squared) values in the early HSC development lineage-tracing dataset^37^. The threshold for isCORAL is set at the maximum F1 score point. Top detected memory genes are highlighted. (D) Memory-gene detection evaluated by AUROC, AUPRC, maximum F1 score, Spearman correlation and Pearson correlation in the HSC lineage-tracing dataset. (E) Scatter plot of ground-truth versus baseline-inferred memory scores in the early HSC development lineage-tracing dataset. Colors and dashed-line conventions are as in (A). (F) ROC and PR curves of different methods for memory-gene detection in the early HSC development lineage-tracing dataset. The point corresponding to the maximum F1 score of isCORAL is highlighted. (G) Exemplary memory-gene detection results across representative methods in the early HSC development lineage-tracing dataset. Violin plots show estimated clonal expression levels from isCORAL, GEMLI, and cell-cycle-regressed SNN, alongside ground-truth clonal expression. Pde4d is a high-memory gene (score = 0.838, rank = 1) despite having dispersed expression in the UMAP space. Both isCORAL (score = 0.343, rank = 34) and GEMLI (score = 0.377, rank = 388) successfully identify it as a high-memory gene, whereas the cell-cycle-regressed SNN substantially underestimates its memory strength (score = 0.021, rank = 3,269) despite its dispersed expression pattern. (H) An additional example highlighting the proliferation-associated gene Top2a in the HSC lineage-tracing dataset. Although highly variable, Top2a exhibits only moderate clonal memory (score = 0.083, rank = 906) and displays clustered expression in the UMAP space. GEMLI (score = 0.766, rank = 6) and cell-cycle-regressed SNN (score = 0.166, rank = 216) assign it much higher memory scores, whereas isCORAL (score = 0.103, rank = 544) only marginally overestimates its memory strength.

**Figure S2.**
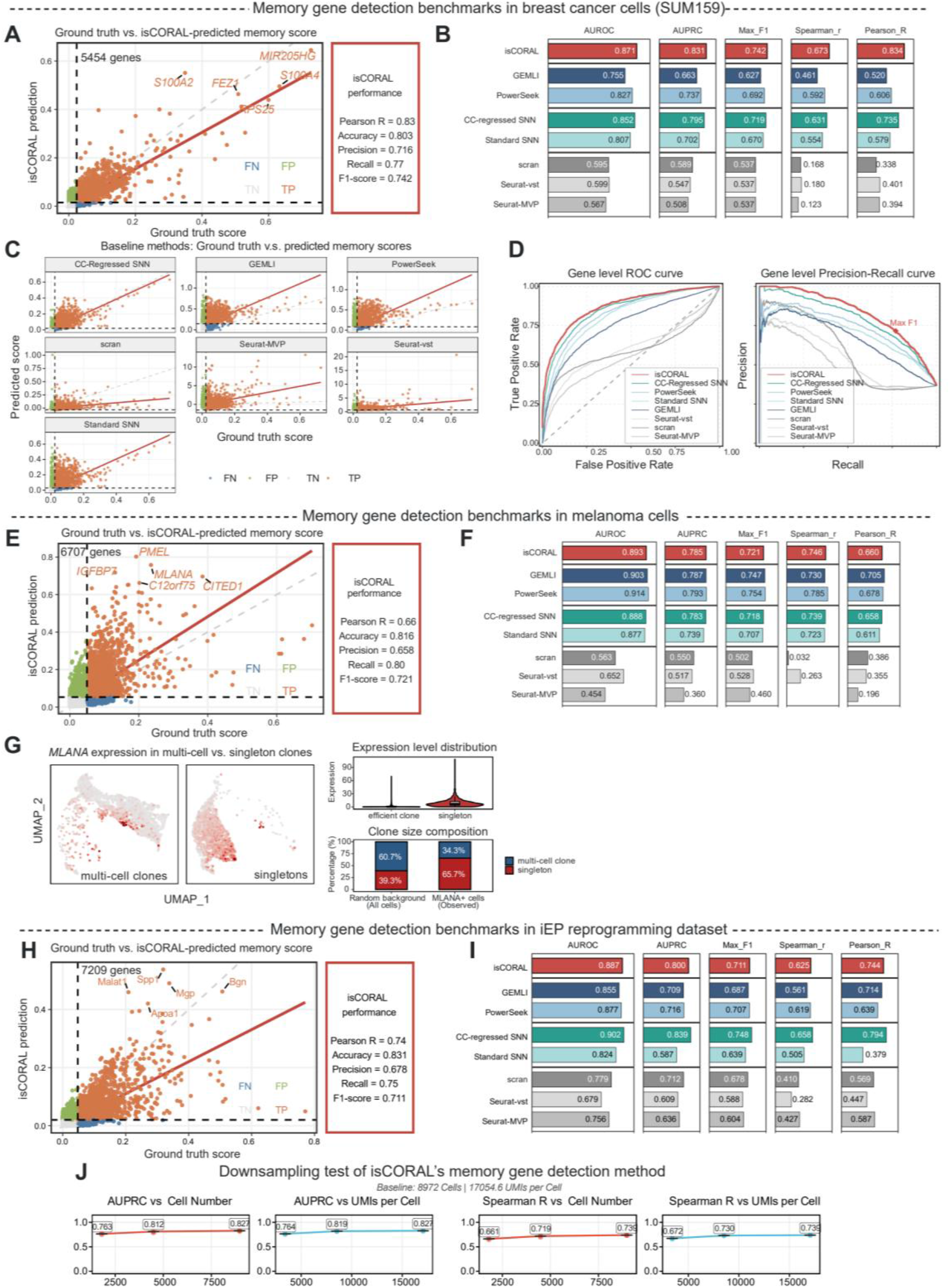
Memory-gene detection benchmarks in SUM159, melanoma and iEP datasets. (A) Scatter plot of ground-truth versus isCORAL-inferred *ω*^2^ (omega-squared) values in the SUM159 lineage-tracing dataset^44^. The majority of top-ranked memory genes (top 2,000) are successfully recovered, with accurate *ω*^2^ quantification (Pearson r = 0.83). The threshold for isCORAL is set at the point maximizing the F1 score. Several top-detected memory genes are highlighted. (B) Comprehensive benchmarking of the memory-gene detection task in the SUM159 lineage-tracing dataset. Evaluated methods include GEMLI, a PCA-based approximation of Power-Seek (Methods), shared nearest neighbor (SNN) baselines (cell-cycle-regressed SNN and standard SNN), and classical highly variable gene (HVG) selection methods (scran, Seurat-Mean-Var-Plot [MVP], and Seurat-Variance Stabilizing Transformation [VST]). Performance is evaluated using recovery metrics (AUROC, AUPRC, and maximum F1 score for the top 2,000 memory genes) and quantification metrics (Pearson and Spearman correlation coefficients). (C) Scatter plot of ground-truth versus inferred memory scores in the SUM159 lineage-tracing dataset. The threshold for isCORAL is set at the point maximizing the F1 score. (D) ROC and PR curves of different methods for memory-gene detection in the SUM159 lineage-tracing dataset. The point corresponding to the maximum F1 score of isCORAL is highlighted. (E) Scatter plot of ground-truth versus isCORAL-inferred *ω*^2^ values in the melanoma fate-mapping lineage-tracing dataset^43^. The threshold for isCORAL is set at the point maximizing the F1 score. Several top-detected memory genes are highlighted. (F) Comprehensive benchmarking of the memory-gene detection task in the melanoma fate-mapping lineage-tracing dataset. Methods and the five benchmark metrics are as in (B). (G) Case study of MLANA in the melanoma fate-mapping lineage-tracing dataset, for which isCORAL inferred higher memory strength than the experimental estimate. (Left) MLANA expression in multi-cell versus singleton clones projected onto the UMAP embedding. (Top right) Quantification of MLANA expression levels in multi-cell and singleton clones. (Bottom right) Proportion of multi-cell versus singleton clone identities among MLANA^⁺^ cells compared to a randomized background. MLANA-high cells are preferentially enriched in singleton clones, a sampling pattern that could contribute to its lower experimentally estimated memory score. (H) Scatter plot of ground-truth versus isCORAL-inferred *ω*^2^ values in the iEP (induced endoderm progenitor) reprogramming lineage-tracing dataset^42^. The threshold for isCORAL is set at the point maximizing the F1 score. Several top-detected memory genes are highlighted. (I) Comprehensive benchmarking of the memory-gene detection task in the iEP reprogramming lineage-tracing dataset. Methods and the five benchmark metrics are as in (B). (J) Robustness of memory-gene detection after downsampling the A549 dataset to 2,500, 5,000 or 7,500 cells, or 5,000, 10,000 or 15,000 UMIs per cell. AUPRC for the top 2,000 ground-truth memory genes and Spearman correlation with the experimentally measured memory scores are shown. Each downsampling condition was repeated three times (Methods); the full dataset contained 8,972 cells with a mean of 17,054.6 UMIs per cell.

**Figure S3.**
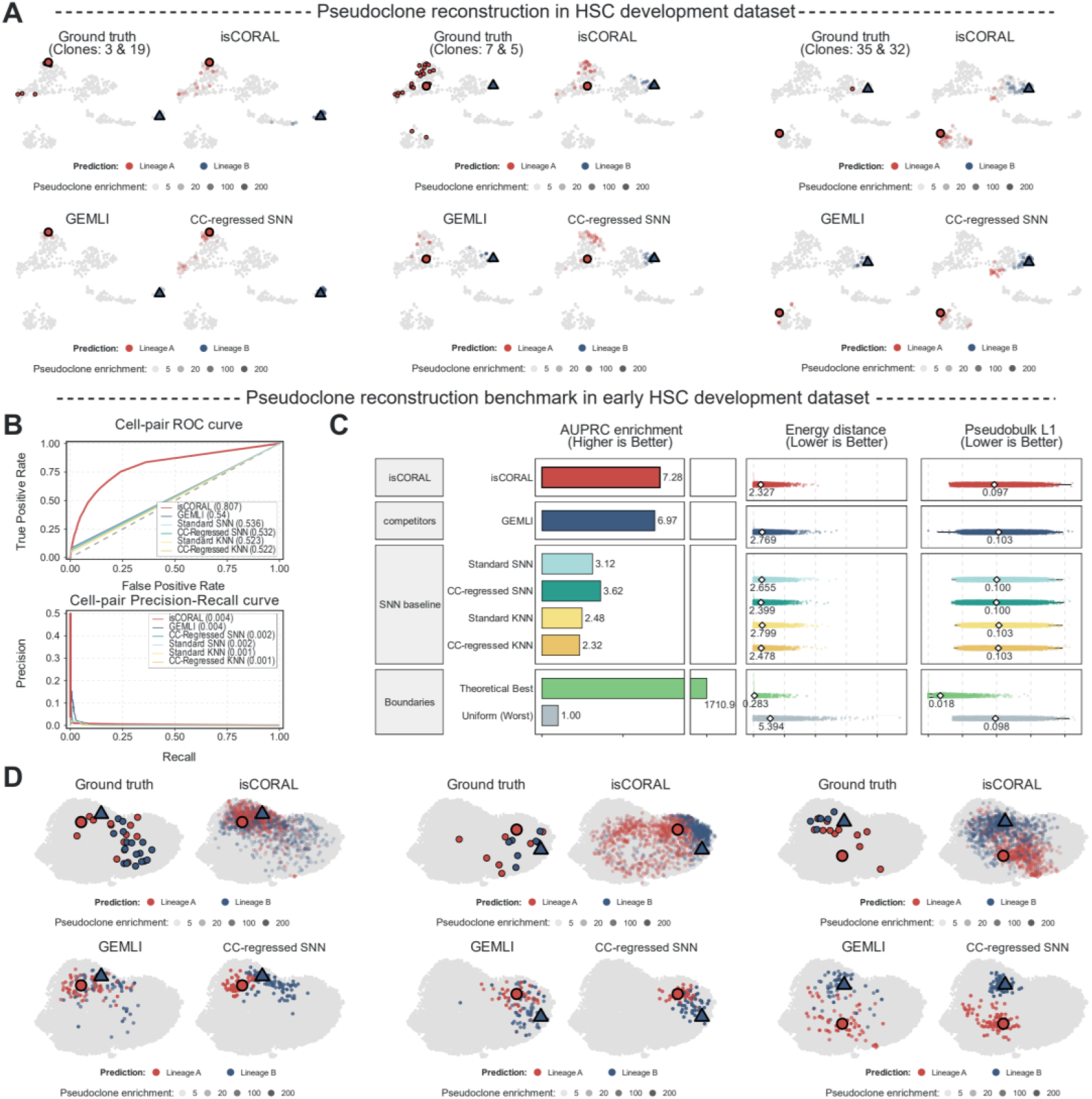
Pseudoclone reconstruction benchmarks in HSC datasets. (A) Additional examples of reconstructed pseudoclones across different methods using the HSC-differentiation lineage-tracing dataset^28^. The outlined red circle and blue triangle denote the seed cells used for reconstruction. Colored dots represent the true or inferred lineages, as described in Figure 3G. (B) ROC and PR curves of different methods for pseudoclone reconstruction on the early HSC development lineage-tracing dataset^37^. Areas under the curves are given in the plot legends. (C) Performance for pseudoclone reconstruction on the early HSC-expansion lineage-tracing dataset. AUPRC enrichment over uniform allocation, energy distance and pseudobulk L1 distance, with display conventions as in Figure 3F. Higher AUPRC enrichment and lower distances indicate better recovery. (D) Examples of reconstructed pseudoclones across different methods using the early HSC-expansion lineage-tracing dataset. The outlined red circle and blue triangle denote the seed cells used for reconstruction. Colored dots represent the true or inferred lineages, as described in Figure 3G.

**Figure S4.**
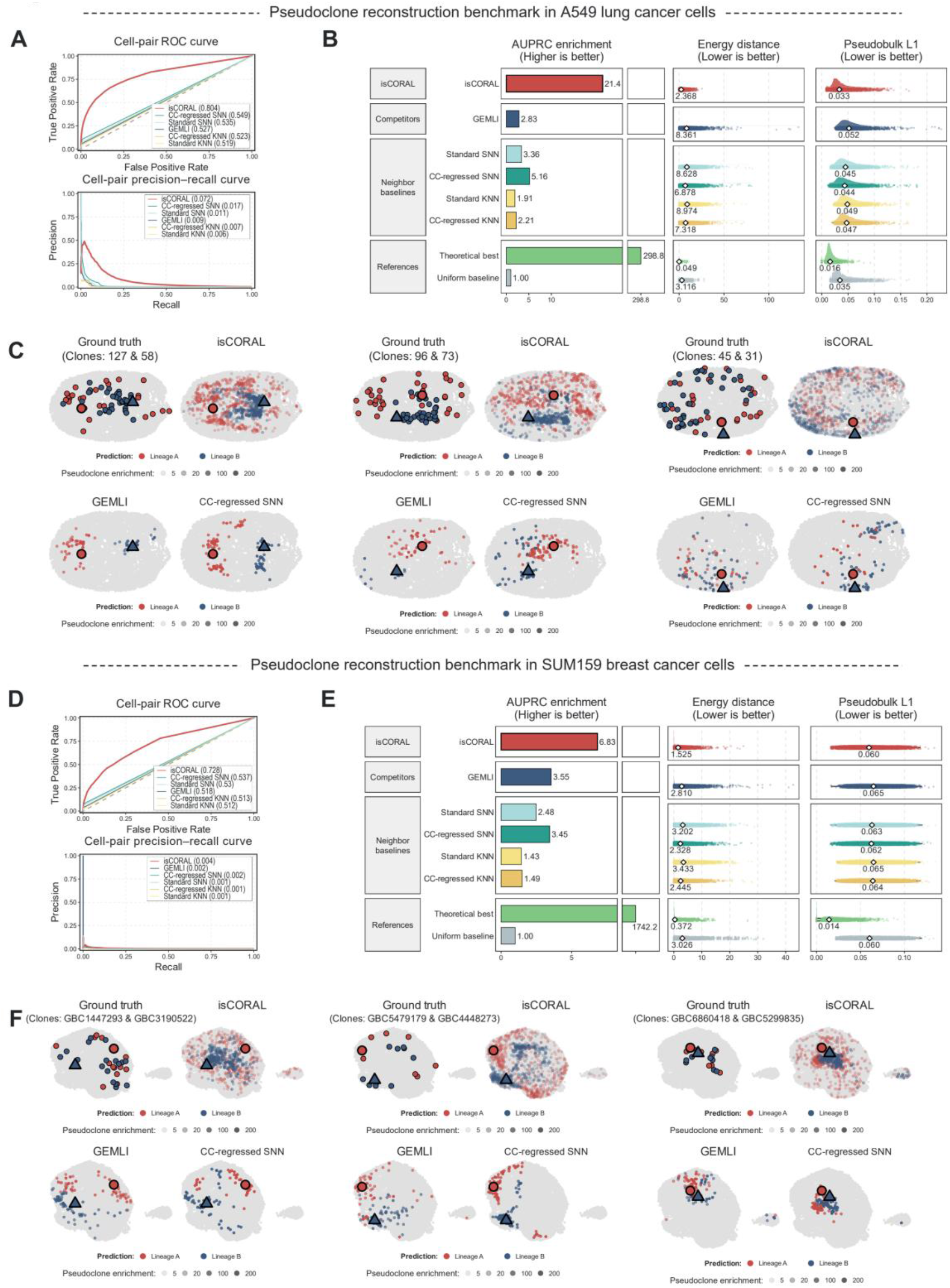
Pseudoclone reconstruction benchmarks in cancer datasets. (A) ROC and PR curves of different methods for pseudoclone reconstruction on the lung cancer A549 lineage-tracing dataset^32^. (B) Performance for pseudoclone reconstruction on the lung cancer A549 lineage-tracing dataset. AUPRC enrichment over uniform allocation, energy distance and pseudobulk L1 distance, with display conventions as in Figure 3F. Higher AUPRC enrichment and lower distances indicate better recovery. (C) Examples of reconstructed pseudoclones across different methods on the lung cancer A549 lineage-tracing dataset. Outlined red circles and blue triangles mark the two seed cells; colors identify their recorded or inferred relatives, with inferred weights shown as enrichment over uniform allocation. (D) ROC and PR curves of different methods for pseudoclone reconstruction on the breast cancer SUM159 lineage-tracing dataset^44^. (E) Performance for pseudoclone reconstruction on the breast cancer SUM159 lineage-tracing dataset. AUPRC enrichment over uniform allocation, energy distance and pseudobulk L1 distance, with display conventions as in Figure 3F. Higher AUPRC enrichment and lower distances indicate better recovery. (F) Examples of reconstructed pseudoclones across different methods on the breast cancer SUM159 lineage-tracing dataset. Outlined red circles and blue triangles mark the two seed cells; colors identify their recorded or inferred relatives, with inferred weights shown as enrichment over uniform allocation.

**Figure S5.**
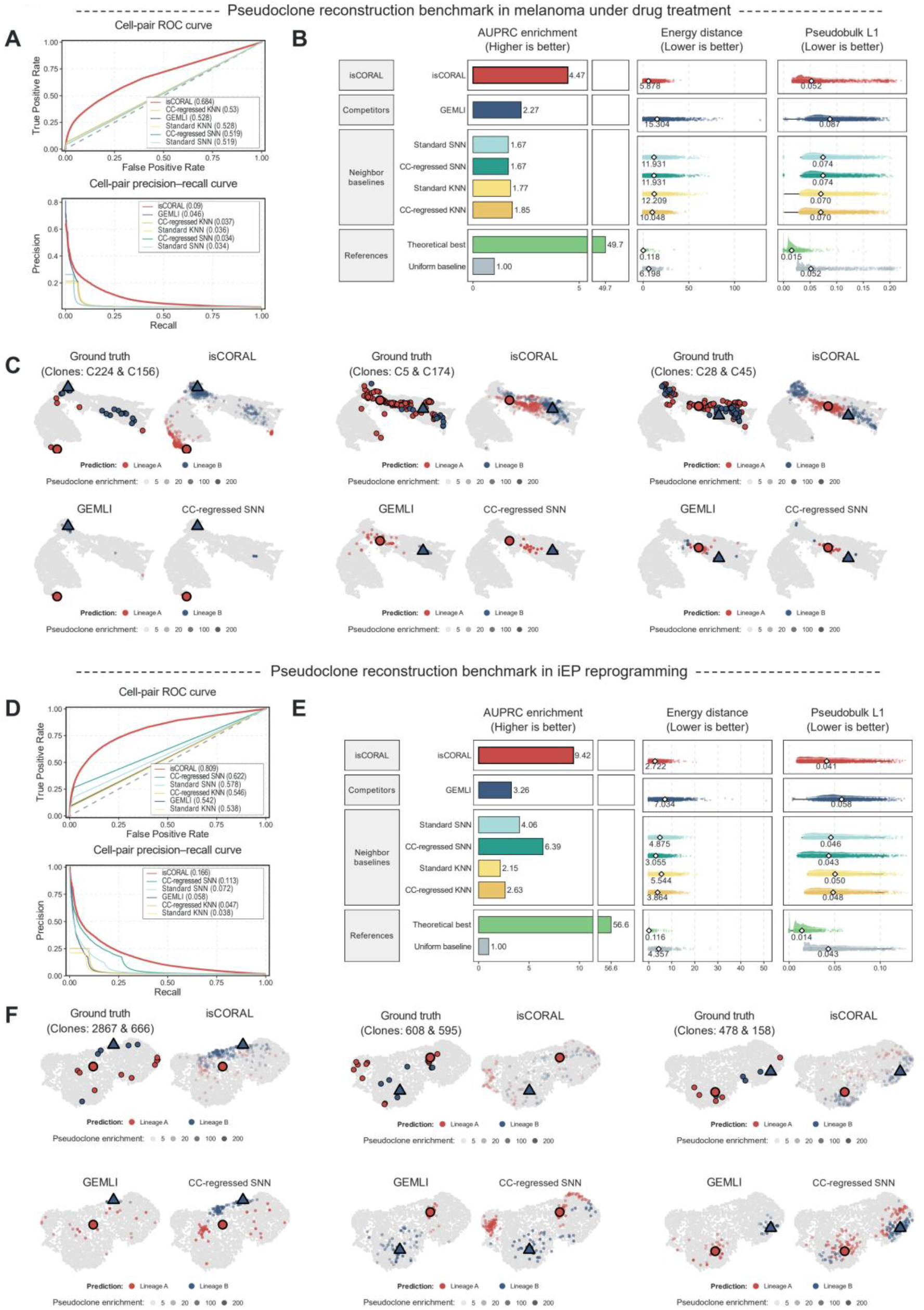
Pseudoclone reconstruction benchmarks in melanoma and reprogramming datasets. (A) ROC and PR curves of different methods for pseudoclone reconstruction on the melanoma lineage-tracing dataset under drug treatment^43^. (B) Performance for pseudoclone reconstruction on the melanoma lineage-tracing dataset under drug treatment. AUPRC enrichment over uniform allocation, energy distance and pseudobulk L1 distance, with display conventions as in Figure 3F. Higher AUPRC enrichment and lower distances indicate better recovery. (C) Examples of reconstructed pseudoclones across different methods on the melanoma lineage-tracing dataset under drug treatment. Outlined red circles and blue triangles mark the two seed cells; colors identify their recorded or inferred relatives, with inferred weights shown as enrichment over uniform allocation. (D) ROC and PR curves of different methods for pseudoclone reconstruction on the iEP reprogramming lineage-tracing dataset^42^. (E) Performance for pseudoclone reconstruction on the iEP reprogramming lineage-tracing dataset. AUPRC enrichment over uniform allocation, energy distance and pseudobulk L1 distance, with display conventions as in Figure 3F. Higher AUPRC enrichment and lower distances indicate better recovery. (F) Examples of reconstructed pseudoclones across different methods on the iEP reprogramming lineage-tracing dataset. Outlined red circles and blue triangles mark the two seed cells; colors identify their recorded or inferred relatives, with inferred weights shown as enrichment over uniform allocation.

**Figure S6.**
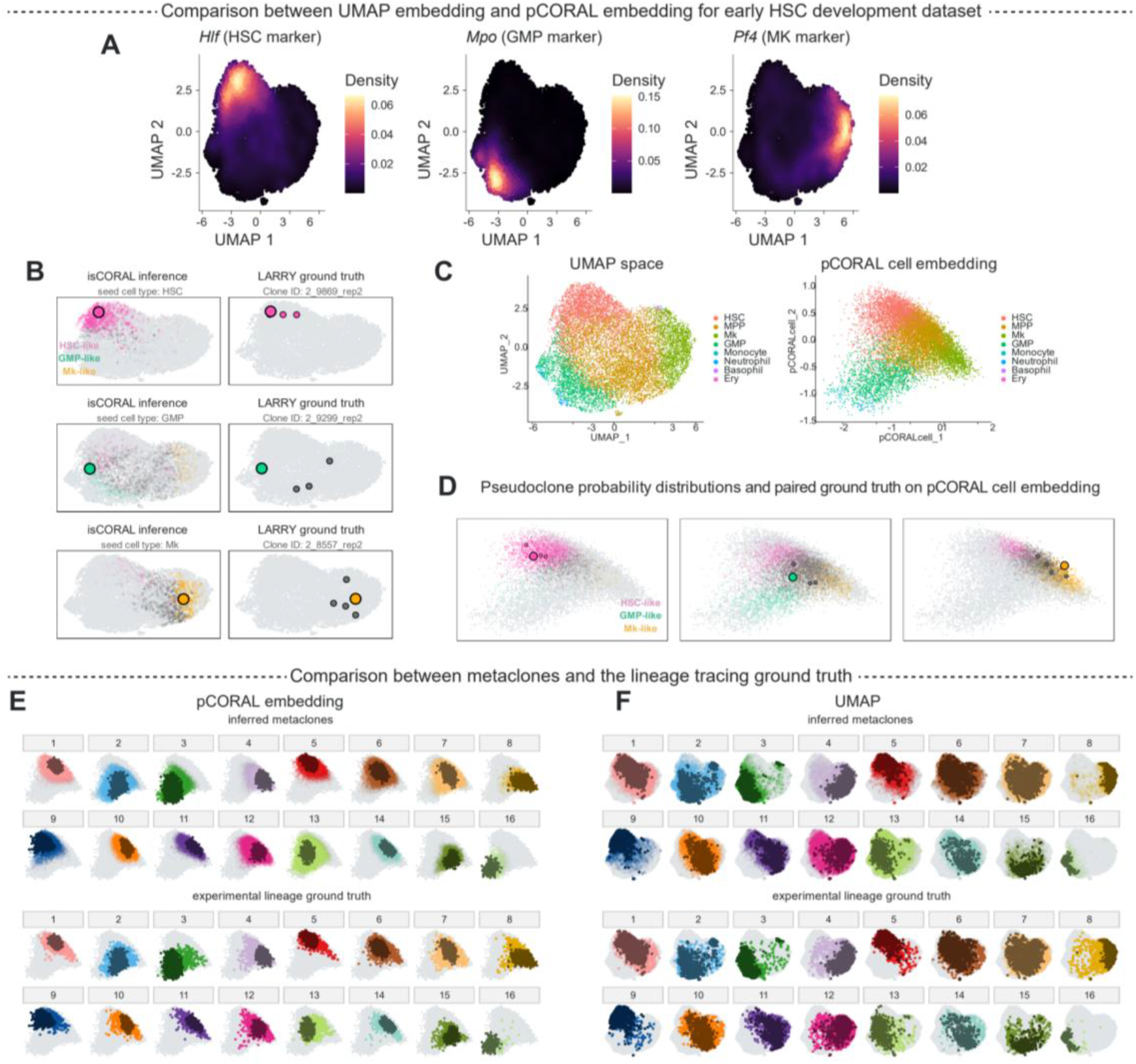
Validation of pCORAL embeddings against LARRY ground truth. (A) Density plots of representative markers during the early HSC-expansion process, visualized in the original UMAP space. *Hlf* (HSC marker), *Mpo* (GMP marker), and *Pf4* (MK marker) are enriched in distinct regions. (B) UMAP visualizations of isCORAL inference results (left) and the paired LARRY lineage-tracing ground truth of representative clones (right). Seed cell types include HSC (top), GMP (middle), and MK (bottom). Cells within the pseudoclone inference or lineage ground truth are highlighted by their cell-type identity, with transparency indicating the inferred pseudoclone weights. The distributions and cell-type proportions of inferred pseudoclones align with the lineage-tracing ground truth. (C) UMAP and pCORAL cell embeddings of the early HSC-expansion process. Cells are colored according to cell-type annotations from the original literature. (D) Merged visualizations of isCORAL inference results and paired LARRY lineage-tracing ground truth of representative clones (same as in B) in the pCORAL cell-embedding space. Seed cell types include HSC (left), GMP (middle), and MK (right). Cells within the pseudoclone inference and lineage ground truth are highlighted by their cell-type identity, with transparency indicating the inferred pseudoclone weights. The distributions and cell-type proportions of inferred pseudoclones align with the lineage-tracing ground truth. (E) (Top) Seed cells and sister cells of inferred HSC metaclones in the pCORAL cell-embedding space. Dark dots represent seed cells; light transparent dots represent sister cells. (Bottom) Experimental lineage-tracing ground truth in the pCORAL cell-embedding space; the real clonal sister cells of the seed cells are plotted as light transparent dots. At the metaclone level, the inferred structure shows high consistency with experimental lineage-tracing results. (F) (Top) Seed cells and sister cells of inferred HSC metaclones in the UMAP space. Dark dots represent seed cells; light transparent dots represent sister cells. (Bottom) Experimental lineage-tracing ground truth; the real clonal sister cells of the seed cells are plotted as light transparent dots.

**Figure S7.**
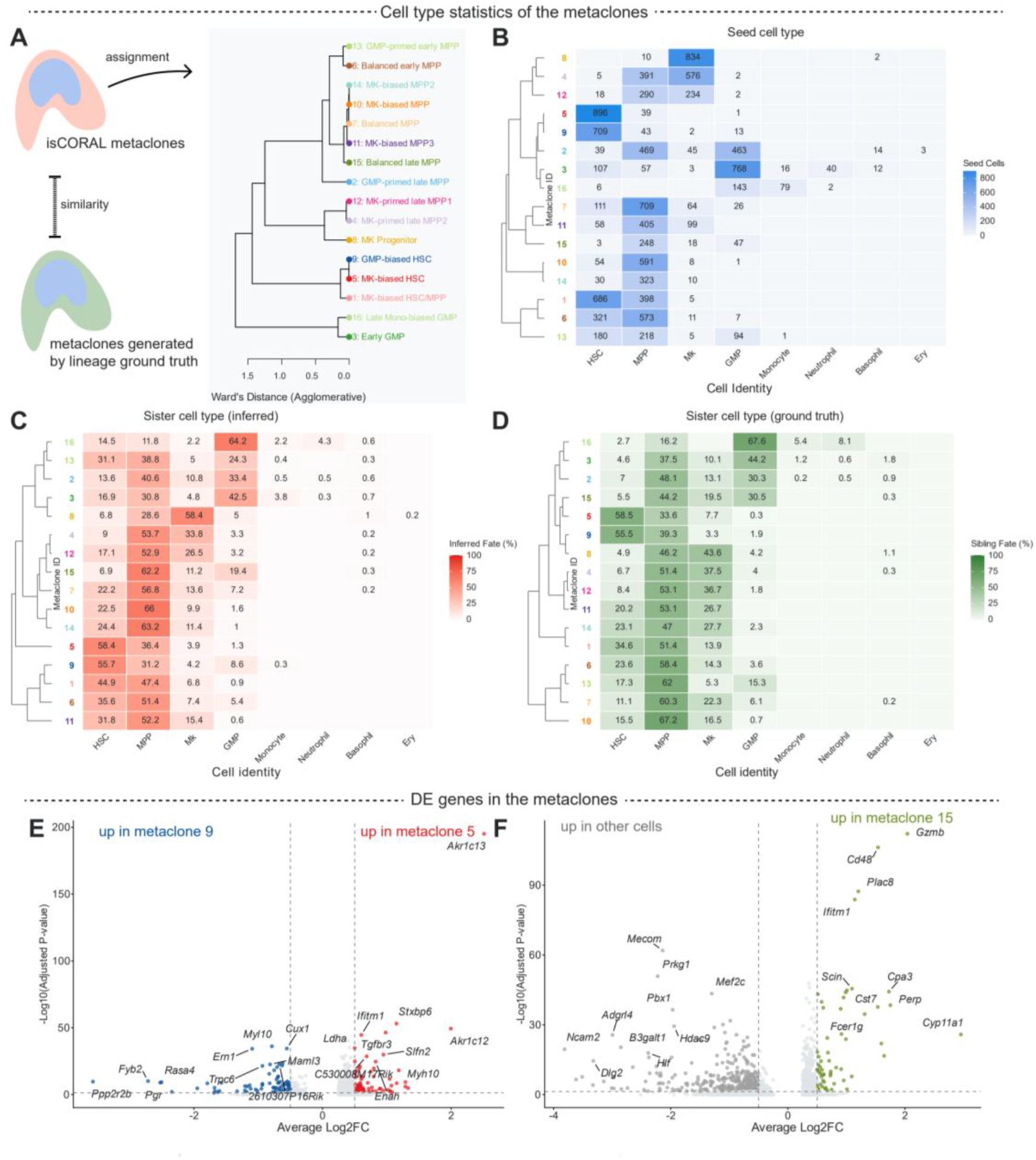
Consistency of metaclone structure with ground truth and the hematopoietic hierarchy. (A) Hierarchical clustering and annotation of the inferred HSC metaclones in Figure 4C. The tree was reconstructed using hierarchical clustering based on the cell-type distributions of the seed cells and sister cells of each metaclone, as shown in (B)–(C). Specifically, the joint matrix of the cell-type distributions of seed cells and inferred sister cells was used for Ward.D2 hierarchical clustering. The dendrogram summarizes similarity in cell-type composition rather than genealogical branching. (B) Seed-cell counts by cell type in each inferred HSC metaclone. Cell types were assigned according to the original literature. (C) Cell-type distribution of the inferred sister cells of each metaclone, weighted by the inferred probability of the sister cells and then normalized. (D) Experimentally measured cell-type distribution of the sister cells of each metaclone. The distribution is normalized so that it can be compared directly with (C). (E) Volcano plot of differentially expressed genes between metaclone 5 and metaclone 9 cells. Each point represents a gene; the x-axis shows the average log2 fold change and the y-axis the −log10 Bonferroni-adjusted P value. Red, significantly upregulated in metaclone 5; blue, significantly upregulated in metaclone 9; gray, not significant (adjusted P < 0.05, |log2FC| > 0.5). Dashed lines mark the significance thresholds; top genes are labeled in italics. Differential expression was tested in seed cells using two-sided Wilcoxon rank-sum tests. (F) Volcano plot comparing metaclone 15 with all other metaclones. Display conventions are the same as in (E): olive, significantly upregulated in metaclone 15; gray, significantly upregulated in all other metaclones; light gray, not significant.

**Figure S8.**
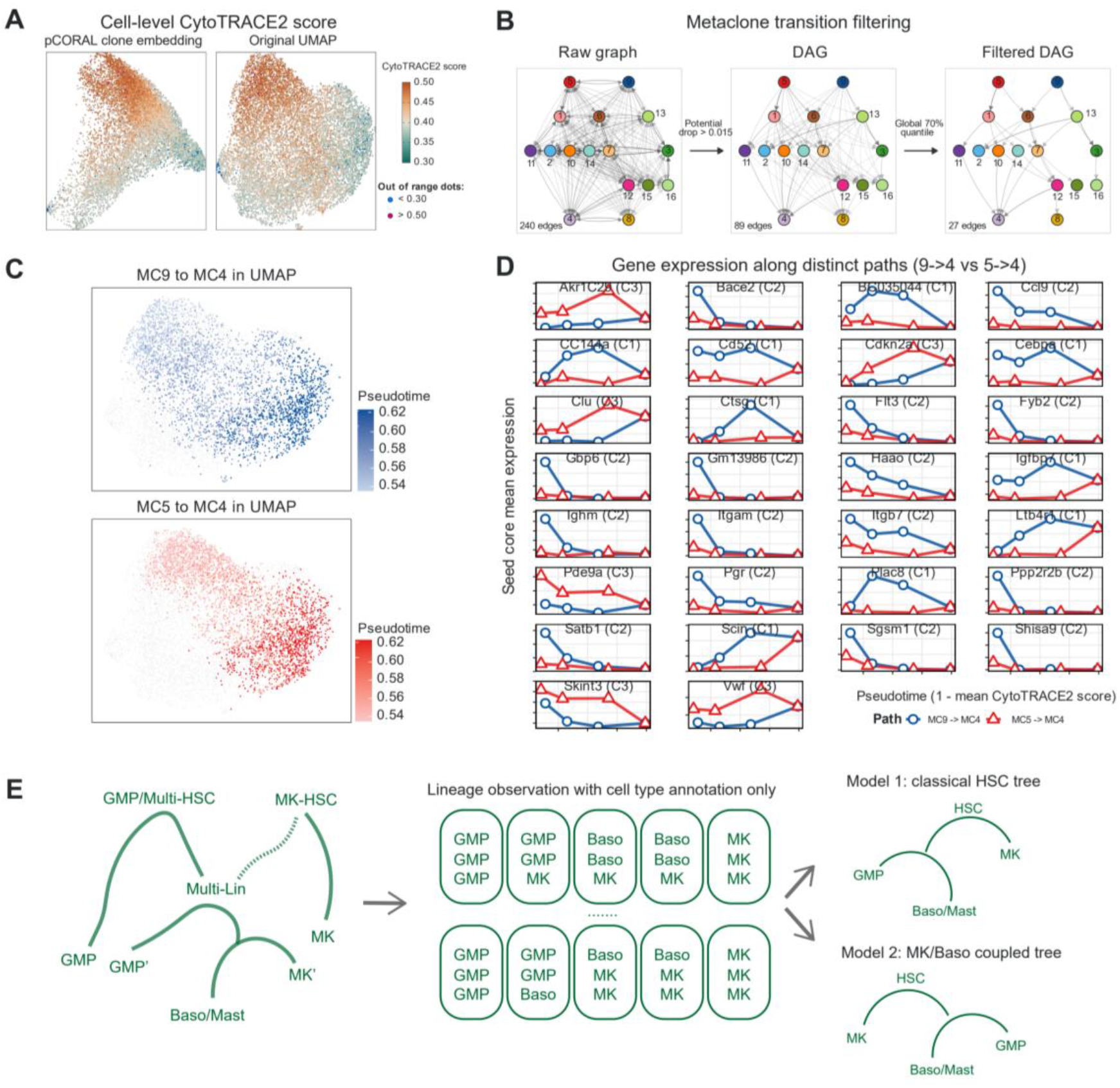
Metaclone transitions and lineage-specific expression trajectories. (A) Cell-level CytoTRACE2 scores in the early HSC-development dataset, visualized in the pCORAL clone embedding and conventional UMAP. Colors transition from teal to orange as the score increases, indicating greater inferred stemness. Blue and magenta points indicate scores below 0.30 and above 0.50, respectively, outside the displayed color range. (B) Filtering of the metaclone transition map. Edges are oriented toward lower CytoTRACE2 developmental potential and retained when the potential drop exceeds 0.015. A global threshold at the 70th percentile of the positive, directionally admissible transition fluxes then retains the strongest 30% to generate the sparse DAG. Internal residency fluxes are retained separately and are not shown as DAG edges. (C) The metaclone 5-1-11-4 path and the metaclone 9-6-7-4 path in the original UMAP embedding. Colors indicate pseudotime, defined as one minus the mean seed-cell CytoTRACE2 score of each metaclone. (D) Expression trajectories of representative differentially expressed genes in Figure 5G. Red triangles and lines represent the metaclone 5-to-4 path; blue circles and lines represent the metaclone 9-to-4 path. Each point shows mean seed-cell expression within a metaclone, plotted against CytoTRACE2-derived pseudotime. C1, C2 and C3 denote modules 1, 2 and 3 in Figure 5G, respectively. (E) Conceptual comparison of cell-type-level lineage observations and lineage-resolved developmental models. Cell-type annotation alone can make alternative models of basophil/mast coupling difficult to distinguish. The isCORAL representation distinguishes fate-biased HSCs and routes through a multilineage progenitor state (Multi-Lin), as defined in Figure 5H.

**Figure S9.**
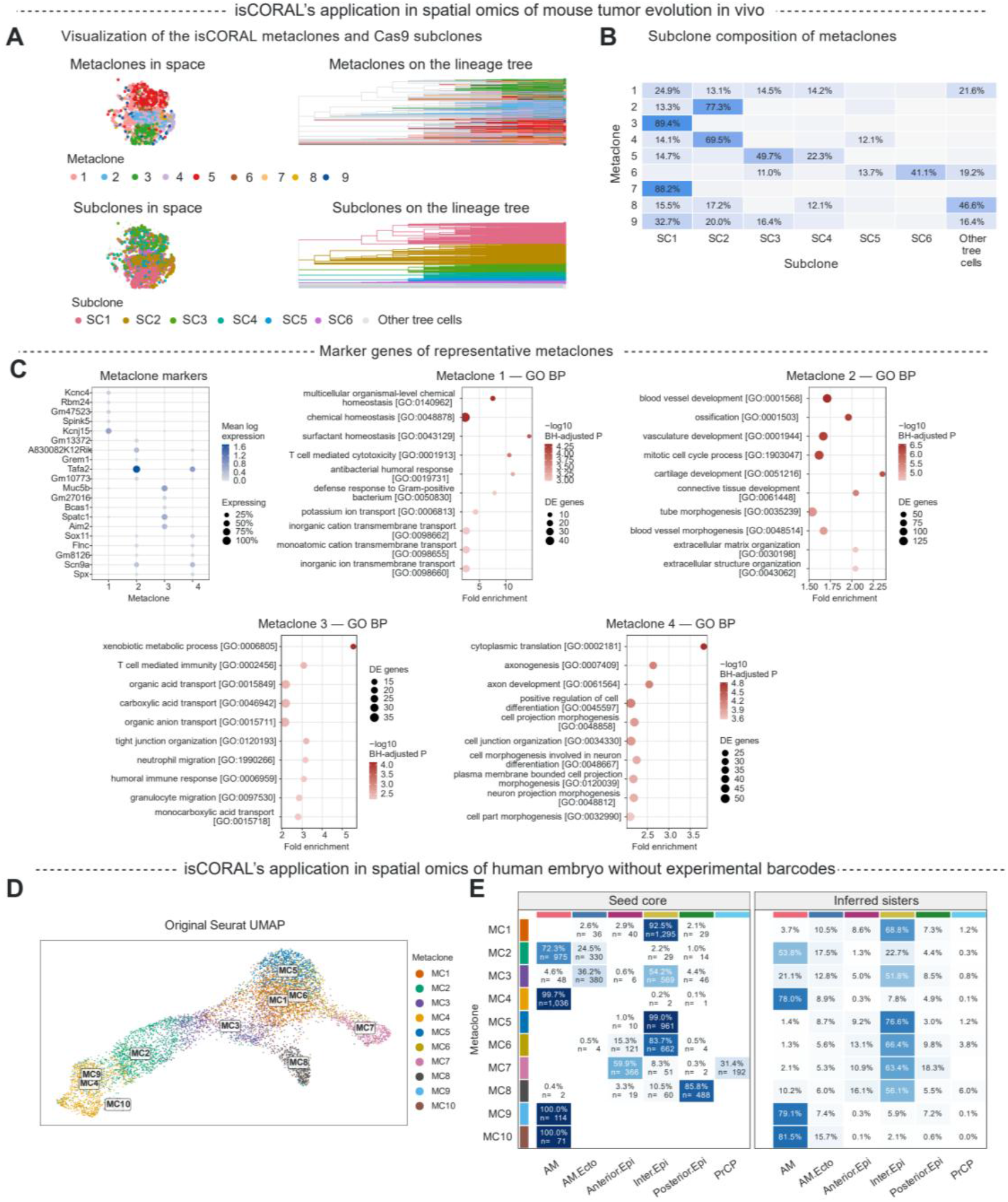
Metaclone analysis of tumor and embryo spatial datasets. (A) Visualization of isCORAL-derived metaclones (top) and Cas9 lineage-tree-derived subclones (bottom) in the original spatial coordinates (left) and on the Cas9 lineage tree (right) of the mouse tumor dataset^38^. (B) Subclone distribution of seed cells in each inferred mouse-tumor metaclone. (C) Marker expression and Gene Ontology (GO) Biological Process enrichment for the four largest mouse-tumor metaclones. Left, the top five differentially expressed genes per metaclone; color indicates mean log expression and dot size the proportion of expressing cells. Right, GO enrichment of differentially expressed genes; the horizontal axis shows fold enrichment, dot size shows the number of contributing genes, and color indicates −log10 of the Benjamini–Hochberg-adjusted P value. (D) Metaclones in the human embryo dataset^39^, visualized in the original UMAP. (E) Cell-type distribution of seed spots and inferred sister spots in each inferred human-embryo metaclone: seed cores are dominated by single annotated cell types, whereas inferred sister distributions span multiple annotated cell types.

**Figure S10.**
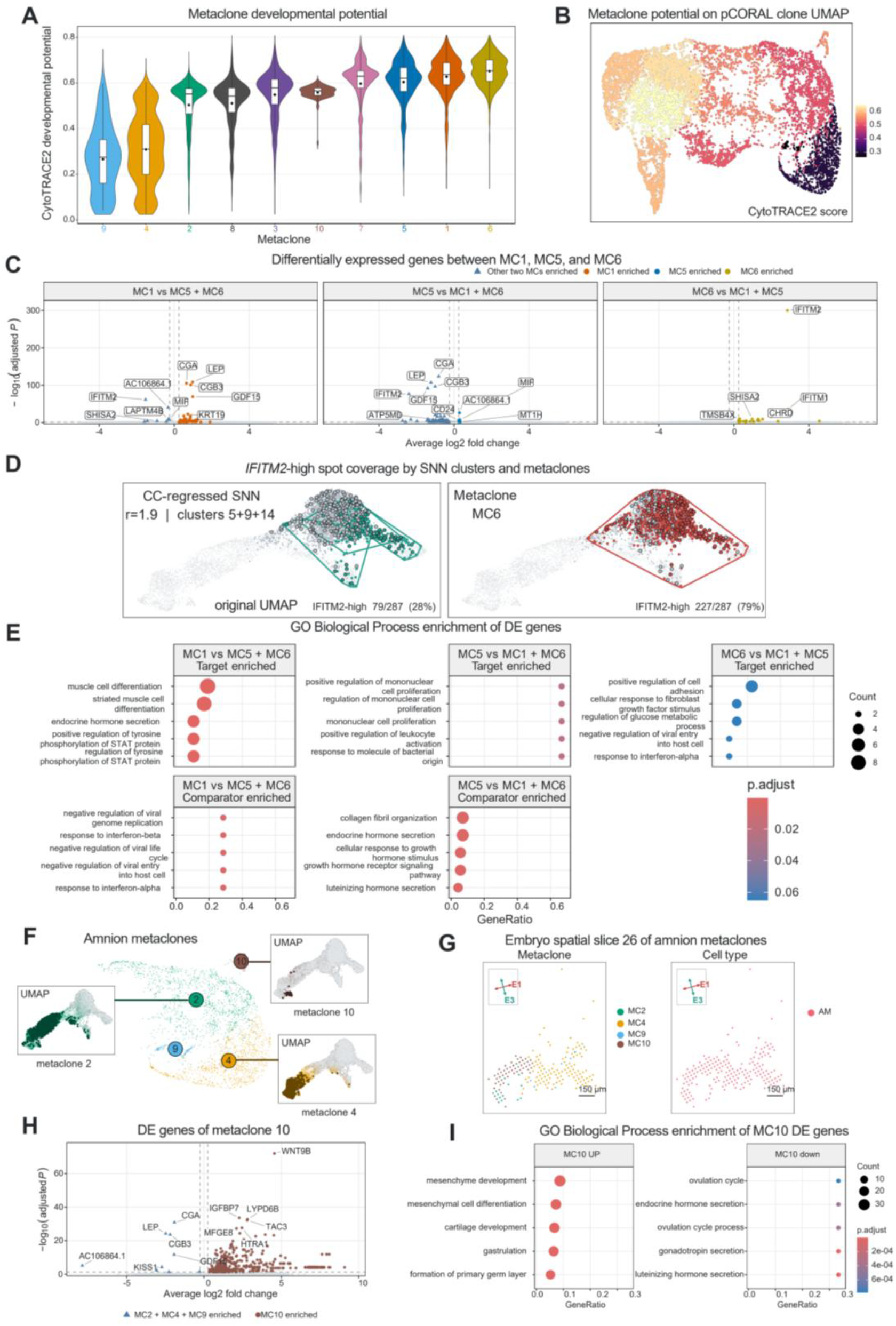
Characterization of human-embryo metaclones. (A) Distribution of CytoTRACE2 scores for each metaclone in the human embryo dataset, sorted from low (differentiated) to high (stem-like). (B) Mean seed-spot CytoTRACE2 scores for each metaclone in the human embryo dataset, visualized in the pCORAL clone UMAP embedding. (C) Differentially expressed genes of metaclones 1, 5 and 6, computed by comparing each target metaclone with the other two pooled metaclones. Each point is a gene; axes show average log2 fold change and −log10 of the Bonferroni-adjusted P value from a two-sided Wilcoxon rank-sum test. (D) Recovery of *IFITM2*-high spots by cell-cycle-regressed SNN clustering and isCORAL metaclones, evaluated within the same pooled MC1/MC5/MC6 population. IFITM2-high spots have expression at or above the 75th percentile among IFITM2-positive spots in this population. For each method, the combination of whole clusters or metaclones maximizing the F1 score was selected; SNN selection was additionally optimized across resolutions (Methods). Selected SNN clusters (left) and metaclones (right) are teal and red. Unselected spots within the comparison population are grey, and spots outside it are light grey. Black outlines mark IFITM2-high spots. The selected SNN clusters recover 79 of 287 IFITM2-high spots (28%), whereas MC6 recovers 227 of 287 (79%). These percentages measure coverage of the IFITM2-high population. (E) Gene Ontology Biological Process enrichment of genes enriched in each target metaclone or its pooled comparator in (C). The horizontal axis shows the gene ratio, dot size shows the number of genes contributing to each term, and color indicates the Benjamini–Hochberg-adjusted P value. (F) The pCORAL clone UMAP embedding and the classical UMAP of amnion metaclones 2, 4, 9 and 10, with conventional UMAP insets for metaclones 2, 4 and 10. In the pCORAL clone UMAP embedding, the colors of circles reflect the metaclone identities. In the UMAP embedding, dark dots highlight seed spots and light dots highlight inferred sister spots. (G) Spatial section 26, enriched for amnion, showing metaclones 2, 4, 9 and 10 (left) and the original cell-type annotation (right). Arrows show the projections of the reconstructed geometric axes E1 (red) and E3 (teal), defined in Figure 6G and Methods. Scale bars, 150 µm. (H) Differentially expressed genes in metaclone 10 compared with pooled metaclones 2, 4 and 9. The horizontal axis shows average log2 fold change; the vertical axis shows −log10 of the Bonferroni-adjusted P value. Differential expression was tested in seed spots using a two-sided Wilcoxon rank-sum test. (I) Gene Ontology Biological Process enrichment of genes enriched in metaclone 10 (left) or in its pooled comparator (right), as identified in (H). The horizontal axis shows the gene ratio, dot size shows the number of contributing genes, and color indicates the Benjamini–Hochberg-adjusted P value.

**Supplementary Table 1.**
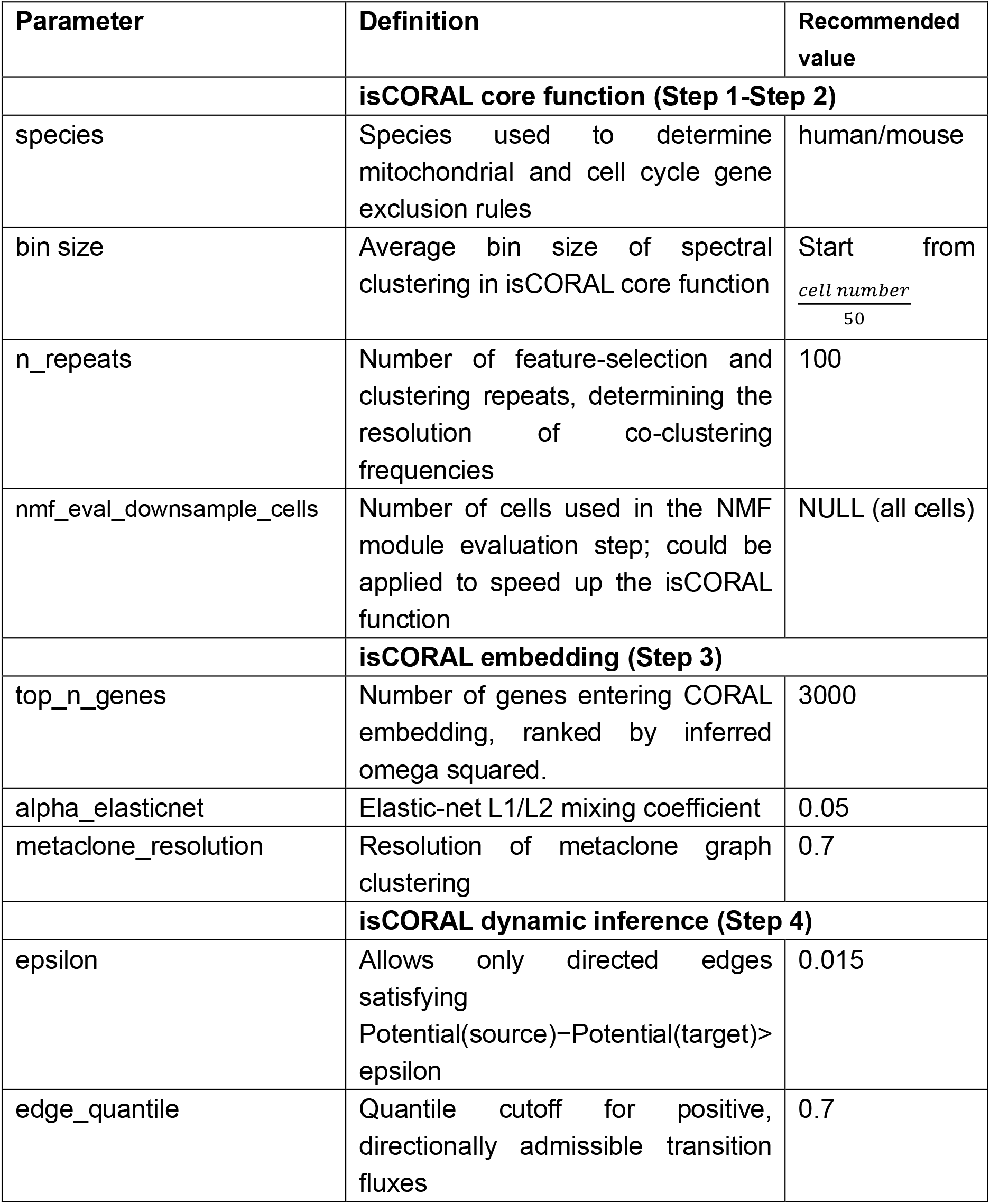
Key tunable parameters of isCORAL.

**Supplementary Table 2.**
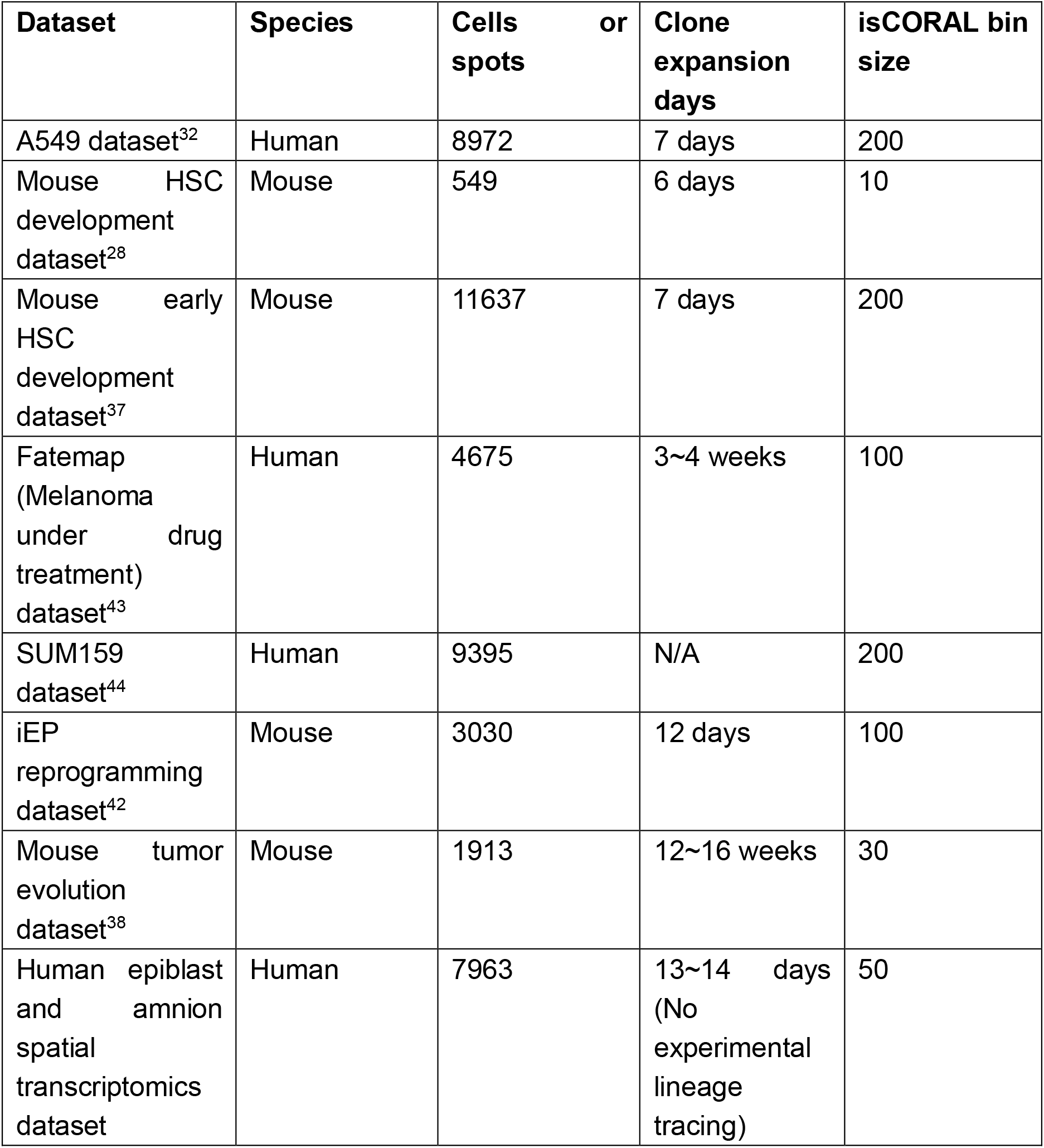
Overview of datasets.

## Methods

### Core task definition of isCORAL

The core task of isCORAL is to infer clonal relationships between cells. Consider an initial cell population labeled with unique heritable clone barcodes and cultured for *d* days before single-cell RNA-seq sampling. Each cell would carry the barcode of its progenitor cell.

For in silico lineage tracing, we seek to estimate whether two cells *i, j* descend from a shared progenitor d days earlier.

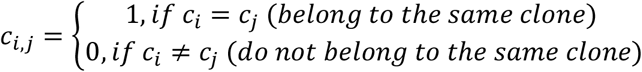

For transcriptome-based lineage inference, the measured expression profile (*M*_*i*_) is affected by measurement noise and biological variation. Transcriptomes therefore do not determine lineage relatedness exactly.

Thus, a practical goal of in silico lineage tracing is to predict the cell–cell clonal-relationship graph:

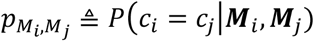

to approximate *c*_*i,j*_.

For one cell with transcriptome *M*_*i*_ in the single-cell RNA-seq dataset of {***M***_*i*_, *i* ∈ *D*}, we define the following probability distribution:

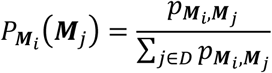

As it approximates the transcriptome distribution of the clone identity:

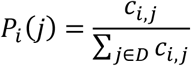

We define 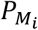 as the pseudoclone distribution of the cell *i*.

### Theoretical motivation of isCORAL’s feature selection strategy

In the single-cell dataset {***M***_*i*_, *i* ∈ *D*}, the measurement of each cell can be represented as ***M***_*i*_ = (*m*_*g*_), *g* ∈ *G*. To simplify the estimation of 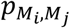, we would like to define a minimal set of key memory genes {*kmg*} ⊂ {*m*}, so that

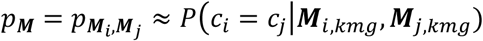

To achieve this goal, we proposed several heuristic rules for selecting memory genes:

*Mean-variance rule: P*(*m*_*i*_ ∈ *KMG*) > *P*(*m*_*j*_ ∈ *KMG*) if gene *i* belongs to a subset of high variance and/or high mean expression while gene *j* is randomly selected. Inspired by the previous publication, the subset can be written as:

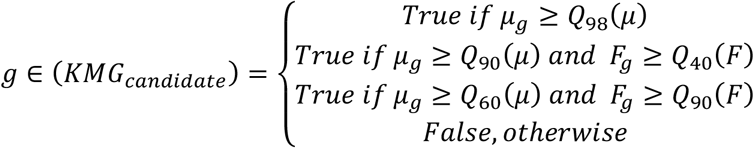

where the *µ*_*g*_ is the mean expression of a gene, and *F*_*g*_ is its Fano factor.

#### Module rule

When two genes *g*_1_, *g*_2_ are very highly correlated, their joint inclusion in the *KMG*: {*kmg*} would be suboptimal. That is:

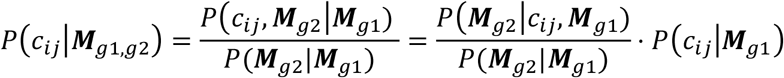

If the *g*_1_ and *g*_2_ are strongly linearly correlated and *g*_1_ is already included in the *KMG*:

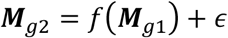

where *E*[*ϵ* ∣ ***M***_*g*1_] = 0 and *Var*(*ϵ* ∣ ***M***_*g*1_) = *σ*^2^ → 0 and

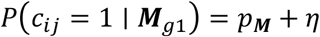

Therefore, we have:

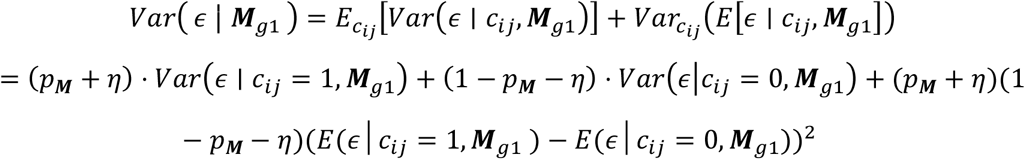

To minimize *Var*(*ϵ* ∣ ***M***_*g*1_) he *Var*(*ϵ* ∣ *c*_*ij*_ = 1, ***M***_*g*1_) *Var*(*ϵ*|*c*_*ij*_ = 0, ***M***_*g*1_) and *E*(*ϵ*│*c*_*ij*_ = 1, ***M***_*g*1_) − *E*(*ϵ*│*c*_*ij*_ = 0, ***M***_*g*1_) must be small.

Meanwhile, we have:

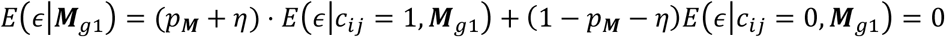

Thus, we expect both *E*(*ϵ*|*c*_*ij*_ = 1, ***M***_*g*1_) and *E*(*ϵ*|*c*_*ij*_ = 0, ***M***_*g*1_) to be approximately zero.

At the resolution of the measurement,

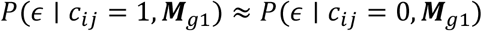

And the *P*(***M***_*g*2_|*c*_*ij*_, ***M***_*g*1_) ≈ *P*(***M***_*g*2_|***M***_*g*1_), *P*(*c*_*ij*_|***M***_*g*1,*g*2_) ≈ *P*(*c*_*ij*_|***M***_*g*1_)

#### Clone bin rule

Guided by these three principles, we developed the first core module of isCORAL, the key feature estimator, to identify a run-specific key feature gene set, KMG_final,q_, from the transcriptomic profiles {M_i_: i ∈ D}.

Let *C* denote true clone identity, *B* the quantile-bin assignment defined by seed gene *g*_1_, and 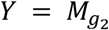 the expression of a candidate gene. We decompose target-gene expression as

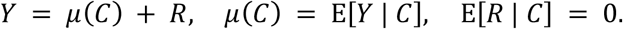

Here, *µ*(*C*) is the clone-specific mean, and *R* is the deviation of an individual cell from its clone mean. The mean expression in bin *k* is

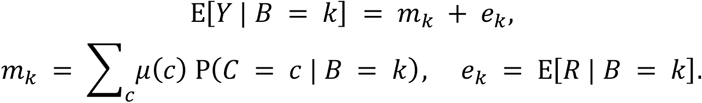

The term *m*_*k*_ reflects differences in clone composition, whereas *e*_*k*_ reflects the association between bin assignment and expression deviations within clones.

Define

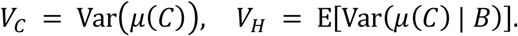

The law of total variance gives

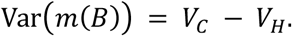

To distinguish population effects from sample statistics, let *θ*_*C*_ and *θ*_*B*_ denote the population proportions of expression variance explained by clone and bin membership, respectively. Assume 0 < Var(*Y*) < ∞.

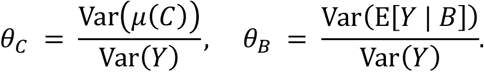

Because E[*Y* | *B*] = *m*(*B*) + *e*(*B*), the exact population relationship is

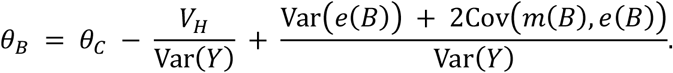

Under the additional assumption that within-clone expression deviations are negligible within each bin, *e*(*B*) → 0.

Thus, a large *θ*_*B*_ supports a large *θ*_*C*_ ndicating lone-associated expression differences in the candidate gene.

### Key memory gene selection

Guided by these three principles, we developed the first core module of isCORAL, the key feature estimator, to identify a run-specific key feature gene set, *KMG*_final,*q*_, from the transcriptomic profiles {*M*_*i*_: *i* ∈ *D*}.

For datasets containing raw counts, we log-normalized the RNA expression matrix using Seurat’s default NormalizeData function. Datasets that lacked a counts layer, such as the iEP reprogramming dataset (CellTag dataset), were analyzed directly from their pre-normalized RNA/data layer without repeated normalization. For each gene *g*, we calculated its mean expression *µ*_*g*_ and Fano factor *F*_*g*_, defined as the population variance divided by the mean. A gene was included in the candidate gene set *KMG*_candidate_ if it satisfied at least one of the following criteria:

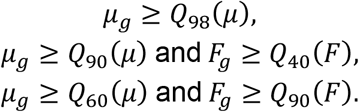

This step enriched the initial pool for potentially informative features.

Cell-cycle genes, identified from Seurat’s cc.genes list, and mitochondrial genes, identified by the MT-prefix in human datasets or the mt-prefix in mouse datasets, were then removed, yielding the cleaned candidate set *KMG*_candidate,clean_.

The expression matrix restricted to *KMG*_candidate,clean_ was subjected to cell-wise L2 normalization and decomposed using RcppML::nmf^69^ with 50 components. One NMF fit was performed per consensus run, with no L1 or L2 regularization applied. For each component, the top *p* = 5 genes ranked by NMF loading were selected and used to generate a component-specific temporary cell partition, *B*_1st_, through Laplace-kernel spectral clustering^70^. The target number of temporary bins was determined by the dataset-specific bin-size parameter:

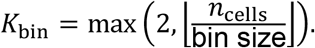

For each component-specific partition, the omega-squared statistic was calculated for every gene. Among genes with mean normalized expression *µ*_*g*_ > 0.05 mega-squared values were ranked in descending order. The validation score of each component was defined as the average omega-squared rank across genes belonging to the intersection of this expression-filtered set and *KMG*_candidate,clean_ ower average rank therefore indicated a more informative component.

The 20% of components with the lowest validation scores (10 of the 50 components) were retained. Their top-five genes were merged and deduplicated to produce *KMG*_final,*q*_ for downstream pseudoclone estimation. This module-validation procedure constitutes the principal time-limiting step of isCORAL. Its computational burden can be reduced through optional downsampling strategies at multiple levels, including UMI-depth downsampling, whole-dataset cell downsampling, candidate-gene subsampling, and cell downsampling during module validation. However, none of these strategies were applied in the benchmark analyses.

### Consensus clustering to generate pseudoclones and memory genes

The second core module of isCORAL is the pseudoclone and memory estimator, which operates downstream of the key feature estimator. Because the features selected in any single iteration can be sparse, we employed consensus clustering to obtain a more robust representation of the latent clonal structure.

For each dataset, we conducted *Q* = 100 independent runs. In each run *q*, the NMF-based feature-evaluation procedure was first applied to generate a run-specific key-feature set, *KMG*_final,*q*_. Laplace-kernel spectral clustering was then performed on the expression matrix restricted to these features,

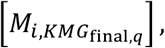

using the same dataset-specific bin-size setting described above. This produced a temporary pseudoclone partition *B*_2nd,*q*_. All 100 runs completed successfully and were included in the subsequent consensus calculation.

The consensus score, or co-association probability, between cells *i* and *j* was defined as the proportion of runs in which the two cells were assigned to the same temporary pseudoclone bin:

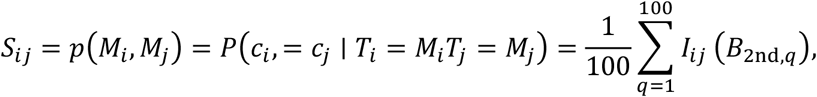

where

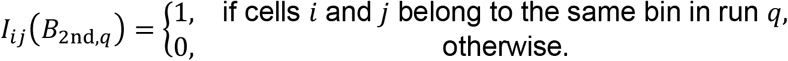

The resulting symmetric co-association matrix *S* captures the stability of the inferred pseudoclone relationships. For downstream probabilistic interpretation and pCORAL embedding, *S* was row-normalized to obtain the pseudoclone probability distribution of cell *i*:

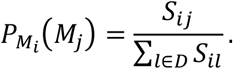

In parallel, each temporary partition *B*_2nd,*q*_ was used to calculate an omega-squared statistic, *ω*^2^, for every gene *g*. The final memory score was obtained by averaging this statistic across all 100 runs and truncating negative values to zero:

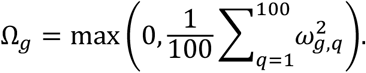

### Pseudoclone-oriented reconstruction of landscape (pCORAL) strategy for pseudoclone and cell embedding

For the visualization and downstream analysis of inferred pseudoclones, we proposed the third module of isCORAL, the pseudoclone-oriented reconstruction of a landscape (pCORAL) strategy to generate robust cell and pseudoclone embeddings within a new CORAL space.

Specifically, we defined a novel cell-cell distance matrix by calculating the squared energy distance^71^ between the inferred pseudoclone probability distributions:

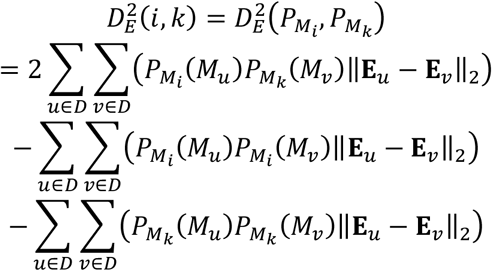

Here, the ‖**E**_*u*_ − **E**_*v*_‖_2_ was calculated on the Seurat-normalized expression matrix of the top *G* memory genes ranked by the consensus memory estimator (e.g., *G* = 3000, without an average-expression filter).

Then, we used this squared distance matrix as the input for classical MDS^72^ to obtain the initial cell embedding **Y**_*MDS*_ (default dimension=30) for each cell.

Next, we applied multivariate elastic-net regression^73^ to project the graph-derived **Y**_*MDS*_ back onto the pure gene expression space. Crucially, we restricted our feature space exclusively to the top *G* memory genes. Let **X** ∈ ℝ^*N*×*G*^ denote this subset matrix after strict cell-wise L2-normalization and gene-wise Z-score scaling.

We sought an optimal linear mapping matrix 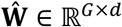. For each target dimension *k*, the projection weights **w**_*k*_ were optimized by minimizing the penalized loss function:

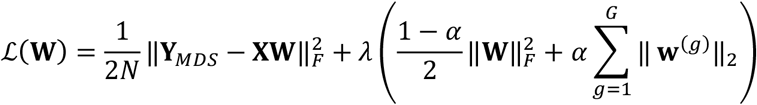

where *α* is the mixing parameter (set to 0.05 to favor Ridge-like preservation of co-expressed modules), and *λ* is tuned via cross-validation. For exceptionally large datasets, a random subsampling strategy was dynamically enabled during this regression.

To account for the coefficient shrinkage induced by regularization, we explicitly recalibrated the variance of each embedded dimension using the scale of the corresponding MDS dimension. The final transcriptomic cell embedding vector, **y**_*cell,k*_, is formally defined as:

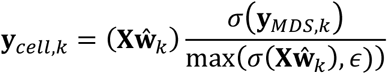

The final cell embedding matrix was then constructed by concatenating these dimensions:

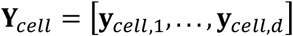

Finally, the pseudoclone embedding, which further mitigates transcriptomic noise, was calculated as the expectation of coordinates under the pseudoclone probability measure:

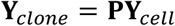

### Dynamic inference module for constructing metaclones

The pseudoclones were further clustered into metaclones in the noise-mitigated clone embedding space **Y**_*clone*_. The shared nearest neighbor (SNN) graph was constructed based on the Euclidean distances across the top dimensions (default: 30) of **Y**_*clone*_. Subsequently, the Leiden algorithm was applied to partition the SNN graph into robust *T* metaclones.

As clusters of pseudoclones, metaclones could be divided into two components: seed cells and inferred sister cells. The seed core of metaclone *t* was defined as:

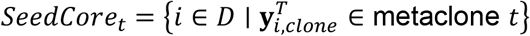

The relevant sister cells were identified as a distribution of combined pseudoclones. To rigorously quantify the outward transition preference excluding self-loops, the one-step outward flux was defined as:

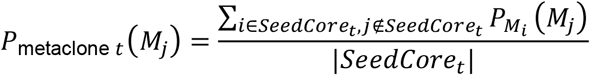

For the dynamic inference module that fits the temporal developmental process, we employed CytoTRACE2^57^ to estimate the average developmental *potential*(*t*) of each metaclone, thereby determining the differentiation direction. Apart from the sister cells, we defined the internal residency flux *R*_*t*_ for metaclone *t* as:

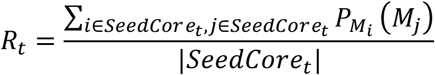

To improve the robustness of the inferred dynamic graph, the directionally admissible off-diagonal fluxes were sparsified before constructing the final transition matrix. Specifically, we defined the raw outward flux from metaclone *t* to *u* as 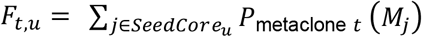. A global filtering threshold *τ* was determined from the positive forward-flux values using a predefined quantile cutoff (default: 0.70) together with a minimum forward-flux cutoff, taking the larger of these two values. Forward fluxes failing to meet the structural threshold *τ* or the temporal potential barrier *ϵ* were penalized to zero, yielding the sparsified forward flux 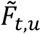:

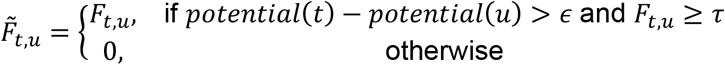

The diagonal residency terms (*R*_*t*_ ere ot ubjected o his iltering. ollowing sparsification, the residency and retained forward-flux terms were jointly row-normalized to obtain the final transition probability *P*_dynamic_(*t* → *u*):

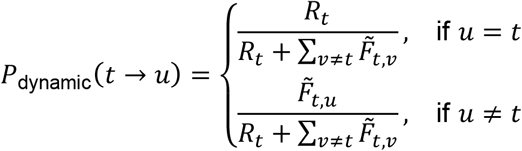

This directionally constrained and structurally sparsified transition matrix *P*_dynamic_ was subsequently utilized to solve the absorbing Markov chain model and estimate terminal fate probabilities.

### Terminal fate, transition flow, and most-probable path inference

Metaclones retaining off-diagonal forward flux were treated as transient states, whereas metaclones satisfying

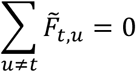

were defined as terminal states. For each terminal metaclone, the entire transition probability was assigned to the diagonal:

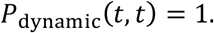

After ordering transient states before terminal states, the transition matrix was written in canonical absorbing form:

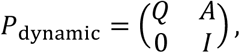

where *Q* contains transitions among transient metaclones and *A* contains transitions from transient to terminal metaclones. The fundamental matrix and terminal absorption-probability matrix were then calculated as

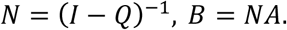

Each element *B*_*a,k*_ represents the probability that a process initiated from transient metaclone *a* is eventually absorbed by terminal metaclone *k*. The expected number of Markov-chain steps before absorption, including diagonal residency steps, was obtained from the row sums of *N*.

Because each retained off-diagonal transition follows a strict decrease in developmental potential, the off-diagonal transition graph is acyclic. A process may persist within a metaclone through diagonal residency transitions but cannot return to that metaclone once it has departed. For any transient source metaclone *a*, which need not be a topological root, *N*_*a,t*_ represents the expected cumulative occupancy of another transient metaclone *t* measured in Markov-chain steps. The probability of reaching *t* from *a* was calculated as

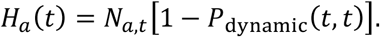

After removing diagonal residency, the state-changing transition probabilities were represented by the jump chain:

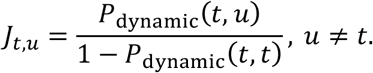

The expected off-diagonal transition flow from source *a* through edge *t* → *u* was defined as

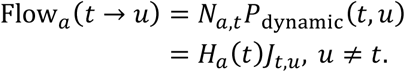

In the acyclic off-diagonal graph, Flow_*a*_(*t* → *u*) equals the probability that the transition *t* → *u* is traversed at least once before absorption. For terminal metaclone *k*, the total incoming flow satisfies

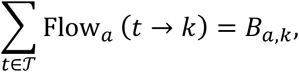

where *T* denotes the set of transient states.

To identify the most probable path from a specified transient metaclone *a* to any reachable downstream metaclone *b*, the path probability under the jump chain was defined as

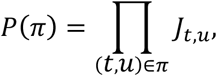

and the most probable path was obtained as

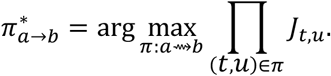

This optimization was implemented as a directed shortest-path calculation by assigning each retained edge the cost *w*_*t,u*_ = − log *J*_*t,u*_.

### Collection and cleaning of single-cell RNA-seq datasets

#### A549 dataset

The preprocessed dataset was downloaded from figshare (https://doi.org/10.6084/m9.figshare.c.8450605). Pre-processing was performed according to the original publications.

#### Mouse HSC development dataset

The preprocessed dataset was downloaded from figshare (https://doi.org/10.6084/m9.figshare.27288630). Pre-processing was performed as described in the benchmark methods below.

#### Mouse early HSC development dataset

The dataset was downloaded from Zenodo (https://doi.org/10.5281/zenodo.14261507). We selected the subset corresponding to “Day-7” and “Female”. We noticed that the original sex annotation was inaccurate, resulting in the inclusion of some male cells. To remove these cells, we calculated a male score based on the average expression of Ddx3y, Uty, Eif2s3y, and Kdm5d. Cells with a male score ≥ 0.2 were filtered out.

For the standard pre-processing of this newly cleaned, single-timepoint dataset, we followed a workflow similar to that of the original publication. Mitochondrial (prefixed with “mt-”) and ribosomal (prefixed with “rps” or “rpl”) genes were explicitly removed from the identified list of highly variable genes. During the scaling process, cell cycle effects were regressed out. Principal component analysis (PCA) was performed on the filtered variable features, and the first 50 PCs were retained. A shared nearest neighbor (SNN) graph was then constructed utilizing the first 50 PCs with a k-parameter set to 20. Finally, UMAP was performed for non-linear dimensionality reduction and 2D visualization using the first 50 PCs and a local neighborhood size of 30.

#### Fatemap (Melanoma drug resistance) dataset

The preprocessed dataset was downloaded from https://www.dropbox.com/sh/my0vgswgrh03li4/AAC9R70cQwixeL-zYGsQumaka?dl=0, and the clone barcode metadata was downloaded from https://github.com/arjunrajlaboratory/ClonoCluster_paper/tree/main/Data/Data_barco des. Pre-processing was performed as described in the benchmark methods below.

#### SUM159 dataset

The preprocessed dataset was downloaded from Zenodo (https://zenodo.org/records/10912157). Pre-processing was performed as described in the benchmark methods below.

#### iEP (induced endoderm progenitor) reprogramming dataset

The preprocessed dataset was obtained from the original data provided in the CoSpar package (https://cospar.readthedocs.io/en/latest/). Pre-processing was performed as described in the benchmark methods below.

#### Mouse tumor evolution dataset

The mouse tumor evolution dataset was retrieved from Zenodo (https://zenodo.org/records/19771805), and the slidetag_1 subset was selected for analysis. Low-quality cells were filtered out by retaining only those with more than 500 and fewer than 5,000 detected genes (nFeature_RNA) and a mitochondrial read fraction of less than 10%. Filtered gene-expression counts were log-normalized to 10,000 counts per cell and scaled using Seurat. The top 2,000 highly variable genes (HVGs) were identified using the variance-stabilizing transformation (VST) method. Principal component analysis (PCA) was performed on the HVGs, followed by graph-based clustering (resolution = 0.5) and uniform manifold approximation and projection (UMAP) dimensional reduction based on the first 15 principal components (PCs). Finally, malignant cells were identified and isolated based on the provided metadata annotation (tumor == TRUE) for downstream analyses.

#### Human epiblast and amnion spatial transcriptomics dataset

The CS6 human pregastrula dataset was retrieved from figshare (https://doi.org/10.6084/m9.figshare.27004099), and the epiblast and amnion subset was extracted based on the original publication. Spatial spots with fewer than 500 or more than 2,500 detected genes were excluded. Gene-expression counts were log-normalized to 10,000 counts per spot and scaled using Seurat. The top 2,000 highly variable genes (HVGs) were identified using the variance-stabilizing transformation (VST) method. Principal component analysis (PCA) was performed on the HVGs, followed by uniform manifold approximation and projection (UMAP) dimensional reduction based on the first 15 principal components (PCs).

### Parameter settings of the benchmark

For both the memory gene detection benchmark and the pseudoclone reconstruction benchmark, we used the default parameters described above. The bin size parameter was determined as in Supplementary Table 2, with a similar 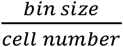 proportion.

### Benchmark of memory gene detection

#### Generation of baselines

Only genes with an average expression level greater than 0.1 in the log-normalized data layer were retained for the memory-gene detection benchmark. The ground-truth memory score was defined as the gene-wise omega-squared effect size from a one-way ANOVA using experimentally observed lineage identities as grouping labels.

In addition to isCORAL, several baseline methods were included for comparison. The Seurat-VST and MVP scores were calculated using Seurat::FindVariableFeatures with the vst and mean.var.plot methods, respectively. Specifically, the standardized variance was used as the Seurat-VST score, whereas the scaled dispersion was used as the MVP score. The Scran score was defined as the biological variance component returned by scran::modelGeneVar.

Because the original Power-Seek^36^ implementation was computationally intensive, we used a faster PCA-based approximation to benchmark the Power-Seek strategy. Briefly, PCA was performed on the log-normalized and scaled expression matrix using the first 30 principal components. The approximate Power-Seek score for gene *g* was calculated as

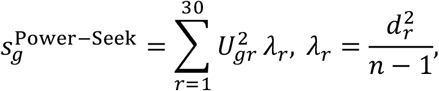

where *U*_*gr*_ is the loading of gene *g* on principal component *r d*_*r*_ is the corresponding singular value, and *λ*_*r*_ is the variance associated with that component. Theoretically, it provides a linear approximation to the original Power-Seek. Because a single gene’s perturbation to the global structure is slight, higher-order terms are expected to be negligible. Consequently, genes with large loadings on high-variance principal components receive higher scores.

Three additional methods (GEMLI^33^, standard SNN clustering, and cell-cycle-regressed SNN clustering) generated inferred cell partitions that were used to estimate gene-wise omega-squared scores. GEMLI was applied as previously described, with a lineage-conversion cutoff of 50. Standard SNN clustering followed the Seurat workflow using the top 2,000 highly variable genes selected by the vst method, the first 30 principal components, and a clustering resolution of 1.5. For the cell-cycle-regressed SNN baseline, S-phase and G2/M-phase scores were additionally regressed out during ScaleData before PCA and clustering.

For both the experimentally observed lineage labels and the inferred partitions generated by these three baseline methods, the gene-wise omega-squared score was calculated as

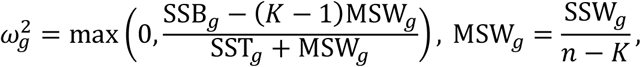

where SSB_*g*_, SSW_*g*_, and SST_*g*_ denote the between-group, within-group, and total sums of squares for gene *g*, respectively; *K* is the number of groups; and *n* is the number of included cells. Experimentally observed lineage identities were used to calculate the ground-truth score, whereas the GEMLI, SNN, and cell-cycle-regressed SNN labels were used to calculate their corresponding baseline scores. Negative or non-finite omega-squared estimates were assigned a value of zero.

#### Metrics

We evaluated the methods using recovery and quantification metrics. Specifically, we calculated the AUROC, AUPRC and maximum F1 score for the top 2,000 ground-truth memory genes as recovery metrics. For the quantitative metrics, we calculated Pearson and Spearman correlations.

### Visualization of example memory genes

To demonstrate the memory gene estimation visually, we plotted the gene expression level at the clone bin levels. For the GEMLI and cell-cycle-regressed SNN methods, we used the clone bins that were used in the omega squared estimation. For the isCORAL method, we applied hierarchical clustering to the (1-consensus score) matrix to obtain approximate discrete clone assignments for visualization.

### Down-sampling test

For the down-sampling test, the cell number or the UMIs per cell were down-sampled before performing the isCORAL algorithm. Each down-sampling was performed three times.

### Benchmark of pseudoclone reconstruction

#### Generation of baselines

For the pseudoclone reconstruction task, the goal is to estimate:

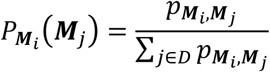

Thus, cell-cell graph reconstruction methods could be benchmarked. GEMLI explicitly provided the confidence matrix, which could be used for benchmarking. For the SNN/KNN baselines, we used standard settings of Seurat, using top 2000 HVGs and top 30 PCA components and extracted the KNN/SNN graph as output. For the cell-cycle-regressed baselines, cell-cycle effects were additionally regressed out during scaling.

To estimate performance boundaries, two empirical control matrices were simulated:

1. Bootstrap Floor: Simulating the empirical upper bound given single-cell sampling limits. We performed random resampling with replacement (bootstrapping) exclusively among cells within the same true clone. To prevent unidirectional sampling artifacts during downstream evaluation, the resulting bootstrap adjacency matrix was strictly symmetrized [(*P* + *P*^*T*^)/2].
2. Uniform Allocation: Simulating a completely uninformative model by assigning equal uniform probabilities across all peer cells.

#### Metrics

We evaluated cell-pair classification and clonal-distribution reconstruction. For the absolute accuracy estimation, we calculated the AUPRC and AUROC of clonally related cell pairs with specific cutoffs in the pseudoclone graph. The AUPRC enrichment (compared with uniform baseline and ground truth baseline) was also reported, especially for those highly imbalanced datasets. For the relative accuracy estimation, we calculated the energy distance in the space of the top 30 principal components between the inferred pseudoclones and the recorded clone distribution for each cell. We also calculated the L1 distance of the inferred pseudobulk expression matrix of pseudoclones and real clones. For the energy distance and the L1 distance matrix, we compared the results with the bootstrap floor and the uniform baseline.

#### Visualization

As the cell number and clone size varied across the datasets, we used a universal FC (fold change) score to visualize the pseudoclone in the benchmark. The FC represented the relative enrichment compared with the uniform allocation matrix and was calculated as:

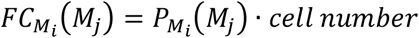

### HSC expansion analysis

For the downstream analysis of the early HSC development dataset, we applied the isCORAL core function and the embedding module with default parameters as described above. The metaclones were identified by Leiden clustering with resolution=0.7.

### Metaclone identity assignments

For each metaclone, the rows of *P*_*norm* corresponding to seed cells were averaged as the ‘sister cells’ distribution, and probability mass directed toward the same “seed” was excluded. To assign and visualize biological relationships among the assigned metaclones, we constructed a joint feature profile by combining the seed-core cell-type composition with the inferred sister-cell fate distribution of each metaclone. The metaclones’ names were assigned manually using the seed cells distribution and the sister cells distribution. Seed-core cell-type counts were row-normalized to percentages before matrix concatenation. Pairwise metaclone dissimilarity was defined from the Pearson correlation of the joint profiles as

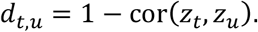

Agglomerative hierarchical clustering was then performed using the Ward.D2 linkage criterion, and the resulting dendrogram was used to organize the metaclones.

### Metaclone fate-bias analysis

For each metaclone, seed-core rows of *P*_norm_ were averaged, and probability mass directed back to the same “seed” was excluded. The remaining masses assigned to MK and GMP cells were normalized within these two cell types, and cell-type bias was calculated as

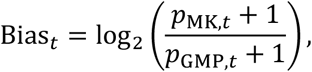

where 1 represents a one-percentage-point pseudocount. Barcode-based bias was calculated using the same formula from MK and GMP cells sharing seed-core barcodes, after excluding the seed-core cells themselves. Positive scores indicate MK bias and negative scores indicate GMP bias. Differences between metaclone 5 and metaclone 9 were evaluated separately for inferred and barcode-based scores using 1,000 two-sided seed-cell label permutations with fixed group sizes and add-one-corrected empirical p values.

Clonal potential to generate both lineages was independently evaluated using experimentally observed lineage barcodes. A clone was classified as possessing MK/GMP dual-lineage potential if it contained at least one MK-lineage cell and one GMP-lineage cell. The proportion of cells belonging to such dual-lineage clones was compared between metaclone 15 and all other metaclones using a two-sided Fisher’s exact test. Together, these analyses quantify both the direct relevant sister-cell bias and the experimentally observed dual-lineage potential of each metaclone.

### Differentially expressed gene analysis of metaclones

Differentially expressed (DE) genes were identified using Seurat::FindMarkers() on the log-normalized RNA expression matrix with a two-sided Wilcoxon rank-sum test. Metaclone 5 was compared directly with metaclone 9, whereas metaclone 15 was compared with all other metaclone-assigned cells. Only genes detected in at least 10% of cells in either group and exhibiting an initial absolute average log_2_ fold change greater than 0.25 were tested. Genes meeting a Bonferroni-adjusted *P* < 0.05 and ∣ avg log_2_FC ∣> 0.5 were deemed differentially expressed. All comparisons were performed on metaclone seed-core cells without incorporating relevant/sister-cell probabilities. Representative DE genes were visualized in metaclone-level feature plots within the CORAL clone embeddings.

### Gene-module analysis of metaclone 15

To characterize stem-associated and lineage-priming transcriptional programs within the metaclone 15 “seed”, module scores were computed from the log-normalized RNA expression matrix using Seurat::AddModuleScore(). Four gene signatures were defined: (i) Baso/Mast-associated progenitor (*Cpa3, Hdc, Gzmb, Gzmc, Cst7, Cd244a, Prg2*, and *Ms4a2*); (ii) HSC-associated stemness (*Ifitm1, Procr, Ly6a, Hoxa9, Meis1, Hlf*, and *Mllt3*); (iii) MK lineage (*Gp5, Vwf, Pf4, Itga2b, Igfbp7*, and *Fli1*); and (iv) granulocytic/GMP lineage (*Mpo, Elane, Ctsg, Lyz2, Cebpa*, and *Csf3r*). Each module score was calculated as the mean expression of the corresponding signature genes minus that of expression-matched control genes.

Module scores were computed for all cells in the metaclone 15 seed core. Two-sided Pearson correlation tests were used to assess single-cell associations between the Baso/Mast score and either the MK-lineage or granulocytic/GMP-lineage score. The HSC-associated stemness score was used to visualize the developmental state of individual cells.

### Metaclone dynamic graph construction in the early HSC development dataset

For the early hematopoietic stem cell (HSC) development dataset, metaclone developmental potential was calculated as the mean CytoTRACE2^57^ score of its seed-core cells. The raw outward flux derived from the aggregated relevant sister-cell distribution was used to define candidate transitions between metaclones. To enforce the expected direction of early HSC differentiation, an off-diagonal transition from source metaclone *t* to target metaclone *u* was considered admissible only when the developmental potential decreased by more than *ϵ* = 0.015. Because every retained transition points toward a sufficiently lower-potential state, the resulting off-diagonal graph is directionally ordered and acyclic.

To reduce weak or potentially unstable connections, the positive directionally admissible fluxes were further sparsified using a global threshold corresponding to their 70th percentile. Fluxes below this threshold were set to zero, yielding the denoised forward-flux matrix *M*_Forward_Clean_. Internal residency fluxes were retained without sparsification because they represent persistence or self-renewal within a metaclone rather than transitions between states.

The retained forward fluxes and residency terms were subsequently combined and row-normalized to construct *P*_dynamic_ etaclones ithout etained off-diagonal outgoing flux were treated as terminal states. A numerical tolerance of 10^−12^ was used during graph construction and near-zero flux determination.

### HSC-specific terminal-flow analysis

For the HSC analysis, metaclone 5 and metaclone 9 were selected as two biologically distinct source metaclones. Metaclone 5 was annotated as an MK-biased HSC state, whereas metaclone 9 was annotated as a GMP-biased HSC state. Source-conditioned expected transition flows were calculated separately as

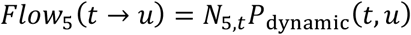

and

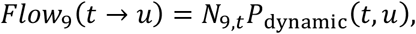

where *N*_*r,t*_ is the expected number of visits to transient metaclone *t* starting from source *r* before absorption.

All terminal metaclones defined by the common denoised transition model were evaluated for both sources. For each source-terminal pair, the terminal absorption probability was obtained from *B*_*a,k*_.

The seed-core identity composition of each terminal metaclone was summarized using the cluster_integrated metadata. Cell identities were grouped into MPP, Mk, GMP, and Others. These proportions described the cellular composition of the terminal “seed”, whereas *B*_*a,k*_ represented the probability of reaching that terminal from metaclone 5 or metaclone 9.

### Comparative analysis of the metaclone 9–metaclone 4 and metaclone 5– metaclone 4 developmental paths

To compare MK-directed development from distinct upstream states, we analyzed paths from metaclone 9 (GMP-biased HSC) and metaclone 5 (MK-biased HSC) to the shared MK-primed terminal state metaclone 4. Diagonal residency probabilities were removed from *P*_dynamic_ nd he emaining ransition robabilities ere row-normalized to construct a jump chain. For each root–terminal pair, the most probable path was identified by maximizing the product of jump-chain edge probabilities, implemented as shortest-path optimization with −log(probability) as the edge cost. The resulting paths were metaclone 9 → metaclone 6 → metaclone 7 → metaclone 4 and metaclone 5 → metaclone 1 → metaclone 11 → metaclone 4.

For transcriptional comparison, log-normalized RNA expression was averaged across the seed-core cells of each metaclone along the two paths. Genes detected in at least 5% of cells in at least one path state were retained. Expression profiles from the two paths were aligned onto a shared CytoTRACE2-derived pseudotime grid, and genes were ranked by their mean absolute inter-path difference relative to their average expression level. The 30 highest-ranking genes were selected for visualization. Their expression trajectories were displayed along both paths and organized by hierarchical clustering using Euclidean distance with complete linkage, yielding three expression-pattern modules.

### Pseudoclone and metaclone inference of the mouse tumor evolution spatial dataset

We applied the isCORAL core function and embedding module to the preprocessed mouse tumor spatial transcriptomics dataset using default parameters, except for a bin size of 30 and metaclone resolution of 0.5.

### Mapping benchmark between pseudoclones and Cas9 lineage tree

The Cas9-derived lineage tree of the mouse tumor evolution dataset was extracted from the Cassiopeia lineage tree reconstruction result in the original research. Only cells retained after preprocessing steps were used for downstream analysis. To benchmark whether the inferred pseudoclones fit the Cas9-derived lineage tree, we first identified the paired branches with the largest F1 recovery as the reference. Then we calculated the energy distance between the pseudoclones and the paired branches in the space of the first 30 principal components, as a measure of the relative reconstruction accuracy. The random control used a ‘random’ pseudoclone assignment, compared with the paired branches as well.

### Differentially expressed gene (DEG) and GO enrichment analysis of metaclones in the mouse tumor evolution spatial dataset

Differential expression (DE) analysis was conducted for the four most abundant metaclones in a one-versus-rest manner. Raw counts were normalized to 10,000 counts per cell and log-transformed (ln(count + 1)). Genes expressed in at least 10% of cells in either group and at least three cells total were assessed using a two-sided Wilcoxon rank-sum test. *P* values were globally Bonferroni-corrected across all features and comparisons. Significant DEGs were defined by an adjusted *P* ≤ 0.05 and | log_2_ FC | ≥ 0.25.

Upregulated and downregulated genes were independently subjected to Gene Ontology (GO) Biological Process enrichment analysis via clusterProfiler^74^ and org.Mm.eg.db. The background universe was defined as all tested genes with valid Entrez ID mappings. Enrichment *P* values were Benjamini–Hochberg-adjusted per metaclone and direction (significance threshold: adjusted *P* ≤ 0.05).

### Pseudoclone and metaclone inference of the CS6 human pregastrula spatial dataset

We applied the isCORAL core function, embedding module and metaclone dynamic inference to the preprocessed CS6 human epiblast and amnion dataset using default parameters, except for a bin size of 50 and metaclone resolution of 0.4. For visualization, we constructed an SNN graph from the pCORAL embedding and applied UMAP using Seurat because the first two MDS coordinates did not clearly separate the trajectories.

### Reconstruction of the geometric axes of the CS6 human embryo

For the spatial distribution of the metaclones, we defined the axes by the following steps: Sections were ordered according to their numerical section identifiers. Within each section, the x and y coordinates were centered on the section centroid and subjected to principal-component analysis without variance scaling. Each section was rotated into its principal-axis coordinate system, with deterministic sign conventions applied to reduce arbitrary orientation reversals between runs. The aligned sections were then stacked along the third dimension at intervals of 20 µm, such that the z coordinate of section *i* was defined as 20(*i* − 1) µm.

For geometric axis estimation and visualization, spatially extreme observations were identified using a robust radial score. Specifically, each coordinate was centered by its median and divided by a robust scale estimate based primarily on the median absolute deviation. The robust radial score was then calculated as the Euclidean norm of the three standardized coordinates. Observations exceeding the larger of a fixed threshold of 6 and an adaptive threshold defined as the median score plus eight robust scale units were omitted from geometric axis fitting and figure generation.

The geometric embryo axes were obtained by eigen decomposition of the covariance matrix of the retained three-dimensional coordinates. Eigenvectors were ordered by decreasing eigenvalue and designated E1, E2 and E3, corresponding respectively to the long, intermediate and short geometric axes of the reconstructed embryo.

### Differentially expressed gene (DEG) and GO enrichment analysis of metaclones in the human embryo dataset

All spatial spots assigned to the predefined embryo metaclones were included in the analysis. MC1, MC5 and MC6 were examined using three one-versus-pooled comparisons: MC1 versus MC5 + MC6, MC5 versus MC1 + MC6, and MC6 versus MC1 + MC5. MC10 was separately compared with pooled MC2, MC4 and MC9 spots. The RNA data layer had previously been generated by normalizing raw counts to 10,000 counts per spot followed by log1p transformation.

Differential expression was performed with Seurat FindMarkers() using a two-sided Wilcoxon rank-sum test. No detection-frequency or fold-change threshold was applied before testing (min.pct = 0, logfc.threshold = 0), both expression directions were retained, and no cells were downsampled. P values were Bonferroni-adjusted by Seurat over the complete RNA feature set within each comparison. Differentially expressed genes were defined by an adjusted P value ≤0.05 and an absolute average log_2_ fold change ≥0.25.

Genes enriched in the target metaclone and genes enriched in the pooled comparator were analyzed separately by Gene Ontology Biological Process overrepresentation analysis using clusterProfiler::enrichGO() and the human annotation database org.Hs.eg.db. Unique HGNC gene symbols passing the differential-expression criteria were used as input, whereas all genes tested in the corresponding comparison with valid gene symbols and finite fold-change estimates constituted the background universe. A direction was analyzed only when at least five eligible genes were available. GO categories containing 10–500 annotated genes were considered, and enrichment P values were adjusted using the Benjamini–Hochberg procedure within each comparison and expression direction. Terms with an adjusted P value ≤0.05 were considered significant.

### Coverage of IFITM2-high spots by cell-cycle-regressed SNN clustering and metaclones

Cell-cycle-regressed shared nearest-neighbor (SNN) clustering was performed on the quality-controlled embryo subset using Seurat. RNA counts were normalized to 10,000 counts per spot and log1p-transformed, and 2,000 highly variable genes were selected using the variance-stabilizing transformation method. S-phase and G2/M-phase scores were calculated using cc.genes.updated.2019 and regressed out during gene scaling. An SNN graph was constructed from the first 15 principal components using 20 nearest neighbours and a pruning threshold of 1/15. Louvain clustering was performed at resolutions from 0.1 to 2.0 in increments of 0.1.

Coverage was evaluated within the same pooled population of MC1, MC5 and MC6 spots for both SNN and metaclone partitions. IFITM2 expression was obtained from the original log-normalized RNA data, without cell-cycle regression. IFITM2-high spots were defined as spots with expression at or above the 75th percentile among IFITM2-positive spots in this pooled population, where positive expression was defined as a value greater than zero.

At each SNN resolution, combinations of whole clusters were evaluated within the comparison population using exact dynamic programming. The same procedure was applied to combinations of the predefined metaclones. For each method, the combination maximizing the F1 score (the harmonic mean of precision and recall for IFITM2-high spots) was selected, with SNN selection additionally optimized across the tested resolutions.

## Data availability

No new experimental data were generated. All datasets analyzed in this study are publicly available from the repositories listed in Methods.

## Code availability

The isCORAL package will be made available at https://github.com/YongjieLin1998/isCORAL_beta. Notebooks reproducing the analyses will be made available at https://github.com/YongjieLin1998/isCORAL_reproduction_notebooks.

## Author contributions

Yongjie Lin designed the project, proposed the algorithm, performed the data analyses, and wrote the manuscript. Yihan Lin designed and supervised the project and wrote the manuscript. All authors reviewed and edited the manuscript.

## Acknowledgments

We thank members of the Lin Lab, especially Yuanzhuo Wang, Yadi Liu, Baoqiang Chen, Jingyuan Lu, Ran Yan, Yifan Xu and Yutong Zhou for helpful discussions regarding analysis. AI-based tools assisted with language editing and manuscript preparation. Yihan L. acknowledges support from the National Natural Science Foundation of China (grant nos. T2325002, 92570207, T2321001, and 32088101).

